# Impact of RNA modification and IL-12 boosts on mRNA-LNP vaccine-elicited CD8+ T cell responses

**DOI:** 10.64898/2026.10.26.761945

**Authors:** Sampa Santra, Michelle Lifton, Benjamin Mildenberg, Georgeanna Morton, Han Tran, Laura Sutherland, Robert Parks, Whitney Edwards, Bette T Korber, Anthony Phan, Christopher Hunter, Garima Dwivedi, Benjamin Davis, Justin Su, Mohamad-Gabriel Alameh, Edward F. Kreider, Susan Pratiksha, Francois Villinger, Tomáš Hanke, Persephone Borrow, Drew Weissman, Barton F. Haynes

## Abstract

Development of effective HIV-1 vaccines requires strategies for inducing both CD8^+^ T cell and neutralizing antibody responses. Nucleoside-modified mRNA-lipid-nanoparticles (LNP) induce more potent antibody responses than unmodified mRNA-LNP vaccines, while unmodified mRNA is typically employed for CD8^+^ T cell-inducing vaccines. Here, we compared the abilities of differing ratios of unmodified versus modified mRNA expressing a SIV conserved-region CD8^+^ T cell immunogen co-administered with IL-12-encoding mRNA-LNP during boosting to induce CD8^+^ T cell and humoral responses in non-human primates, assessing responses in ChAdOx1-primed controls in parallel. Serial priming and boosting with fully-unmodified mRNA elicited higher-magnitude, more CD8-biased T cell responses than fully-modified mRNA, that were durably maintained after the IL-12-encoding mRNA boosts. Inclusion of 30% nucleoside-modified mRNA enhanced antibody titers, suggesting potential benefits to a mixed-composition vaccine.

**One sentence summary:** Varying ratios of modified and unmodified nucleoside containing mRNA-LNP vaccines and addition of IL-12 elicits CD8^+^ T cell responses in monkeys.

## INTRODUCTION

Human immunodeficiency virus type 1 (HIV-1) vaccines will need to elicit both cellular and humoral immunity to combat infection effectively. Although broadly neutralizing antibodies (bnAbs) can prevent HIV-1 acquisition (*1*), eliciting and durably maintaining the high titers of bnAbs needed for protection by vaccination remains elusive. Potent bnAbs are generated in a minority of people living with HIV (PLWH) but typically require elevated viral loads and several years to develop (*2–6*). In contrast, virus-specific CD8^+^ T cell responses do not block the establishment of infection, but do play a key role in viral containment in acute HIV-1/simian immunodeficiency virus (SIV) infections (*7–10*) and sustained viral control in a subset of elite controllers who mount T cell responses to conserved epitopes in Gag and other HIV-1 proteins (*11–14*). Vaccine-elicited CD8^+^ T cell responses are also associated with reduced peak viremia and viral set points in non-human primates (NHPs) (*14–16*) Moreover, a vaccine vectored by the molecular clone 68-1 of rhesus cytomegalovirus (RhCMV68.1) and expressing a simian immunodeficiency virus (SIV) immunogen that elicited atypical major histocompatibility complex (MHC)-E-restricted CD8^+^ T cell responses enabled ∼55% of immunized animals to mediate rapid clearance of SIV following transmission (*17–19*).

A recent NHP study has demonstrated that vaccine-induced T cell responses protected macaques from mucosal infection with simian human immunodeficiency virus (SHIV) in the presence of low titers of serum neutralizing antibodies (*20*). This study used a complex series of viral vectors for the induction of CD8^+^ tissue resident memory T cells (TRM), which were hypothesized to synergize with low serum neutralizing antibody to mediate protection (*20*). While this study requires confirmation, synergy between CD8^+^ T cells and bnAbs in suppressing HIV-1 replication has also been demonstrated *in vitro* (*21*). In therapeutic settings in both SHIV-infected NHPs and PLWH, bnAb administration has also been shown to be associated with expansion of virus-specific CD8^+^ T cells that are associated with long-term viral control (*22–28*).

Nucleoside-modified mRNA lipid nanoparticles (LNP) are a leading platform for many infectious disease vaccines and have been shown to induce strong T_H1_-biased-CD4^+^ T follicular helper cell (T_FH_) and neutralizing antibody responses, together with CD8^+^ T cell responses in mice, NHPs and humans (*29–32*). Notably, the long term protection from severe disease and death due to COVID-19 induced by either the vaccine or infection is mainly mediated by CD8^+^ T cell responses [reviewed in (*33*)]. By contrast, although unmodified mRNA-LNP vaccines drive lower immunogen translation and humoral responses than nucleoside-modified mRNA-LNPs, they stimulate high-level type I interferon (IFN) production and can elicit strong CD8^+^ T cell responses (*34–37*). Unmodified mRNA designs are typically chosen for CD8^+^ T cell induction in cancer vaccine studies (*38*).

Thus, based on these properties of unmodified and modified mRNAs, here investigated whether mRNAs can be designed to elicit high magnitude CD8^+^ T cell responses while maintaining the elicitation of humoral responses against SIV/HIV antigens. We compared the induction of CD8^+^ T cell and humoral responses in rhesus monkeys immunized with different ratios of unmodified vs modified mRNA-LNPs encoding a T cell immunogen, SIVconsv239, an SIV_239_ analog of the most conserved regions of HIV proteome, HIVconsvX (*39*) **(Fig. S1A-C)**. The magnitude and breadth of CD8^+^ T cell responses to peptides spanning the HIVconsvX sequence in PLWH has been shown to correlate with high CD4^+^ T cell counts and low HIV loads (*39*). In people living without HIV (PLWoH) prime-boost vaccination with chimpanzee adenovirus (ChAdOx1) (*40*) and modified vaccinia Ankara (MVA) vectors encoding HIVconsvX elicited broad CD8^+^ T cell responses that mediated cross-clade inhibition of HIV-1 replication *in vitro* (*41–43*). Thus, for comparison with mRNA-LNP, we also immunized a group of monkeys with ChAdOx1.SIVconsv239, as this vaccine has been shown to induce high magnitude CD8^+^ T cell responses (TH, Hansen and Picker, unpublished). To enhance late boosting immunizations, we added an mRNA-LNP adjuvant encoding macaque IL-12 to further increase CD8^+^ T cell responses (*44*) **(Fig S1D)**. We show that mixtures of modified and unmodified mRNA-LNP plus mRNA-LNP encoding IL-12 can induce both anti-HIV CD8^+^ T cell and antibody responses.

## RESULTS

### Immunization with unmodified mRNA elicits higher magnitude and more durable T cell responses than modified mRNA in rhesus monkeys

To compare the induction of CD8^+^ T cell and humoral responses in NHPs immunized with differing ratios of unmodified versus modified mRNA-LNPs encoding a T cell immunogen (SIVconsv239), we immunized four groups of NHPs (n=5/group) intramuscularly with 50 μg of mRNA-SIVconsv239 on weeks 0, 4 and 8 as follows: 100% 1-methylpseudouridine modified mRNA-LNP (group 1), 70% modified and 30% unmodified nucleoside mRNA-LNP (group 2), 30% modified and 70% unmodified mRNA-LNP (group 3) and 100% unmodified mRNA-LNP (group 4). Monkeys in group 5 received ChAdOx1.SIVconsv239 as positive control for CD8^+^ T cell induction. Immunizations of group 5 monkeys started 4 weeks later than mRNA-LNP immunizations, and two immunizations were given on study weeks 4 and 8 **(Table S1).** We analyzed CD8^+^ T cell responses by SIV Gag p11C (*8, 45*) tetramer staining of whole blood of *Mamu-A*01-*expressing monkeys in each group and tested peripheral blood mononuclear cells (PBMC) in ELISpot and intracellular cytokine staining (ICS) assays in all animals using a pool of peptides spanning the SIVconsv239 immunogen sequence. Gating strategies and representative examples of tetramer staining are shown in **Fig. S2**.

Two weeks post-3^rd^ mRNA immunization on week 10, unmodified mRNA (group 4) elicited 24.5-fold higher mean tetramer-reactive CD8^+^ T cell responses than 100% modified mRNA (group 1) immunizations, with responses of intermediate magnitude being observed in the groups receiving 30% and 70% modified mRNA (**Fig. 1A**). By week 16 of the study these *ex vivo* CD8^+^ T cell responses had contracted, although group 4 monkeys maintained 11-fold higher mean responses than group 1 monkeys. The responses in group 4 animals were comparable to those elicited by two priming immunizations with ChAdOx1-vectored SIVconsv239 (**Fig. 1A**).

**Fig. 1.**
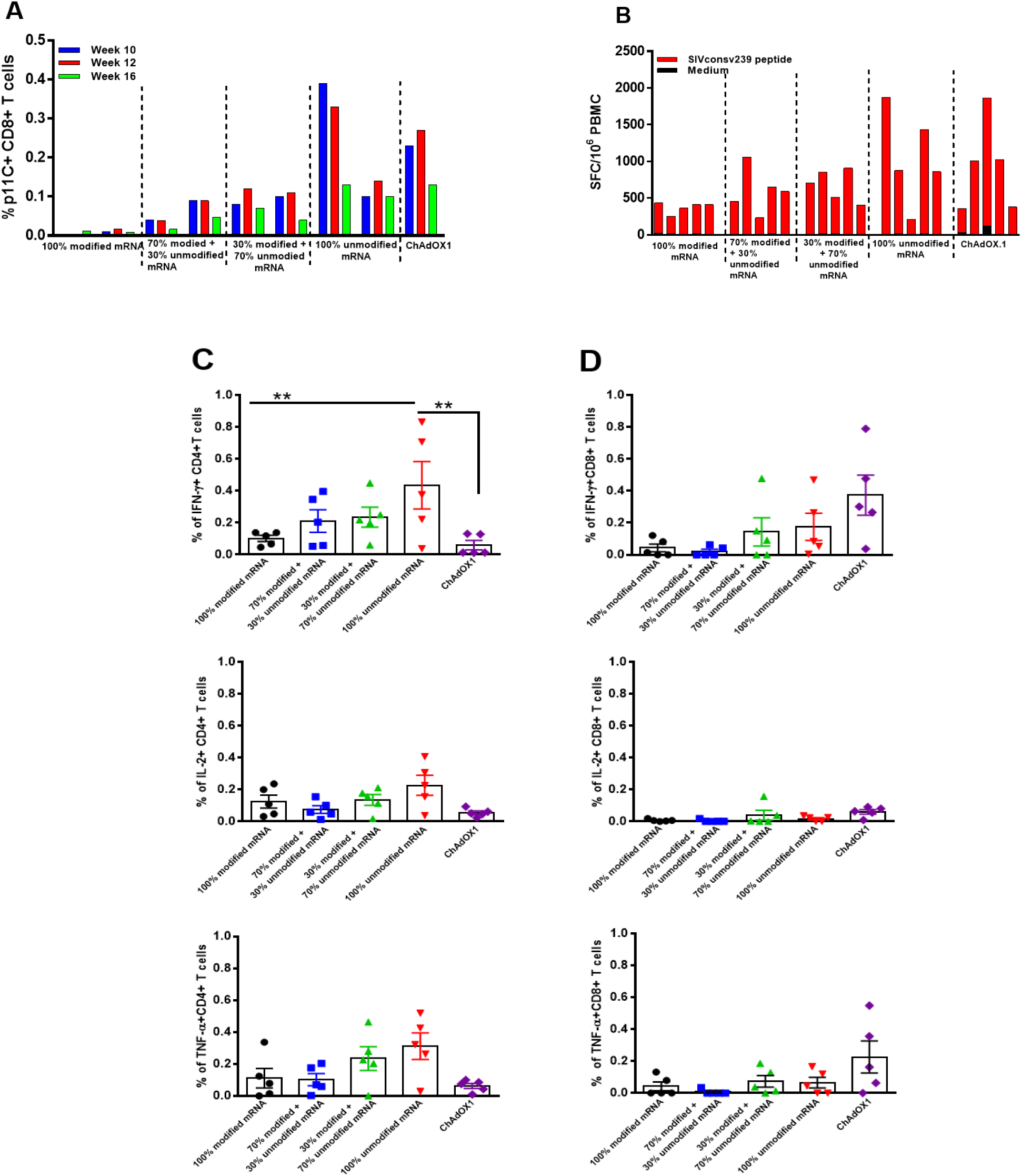
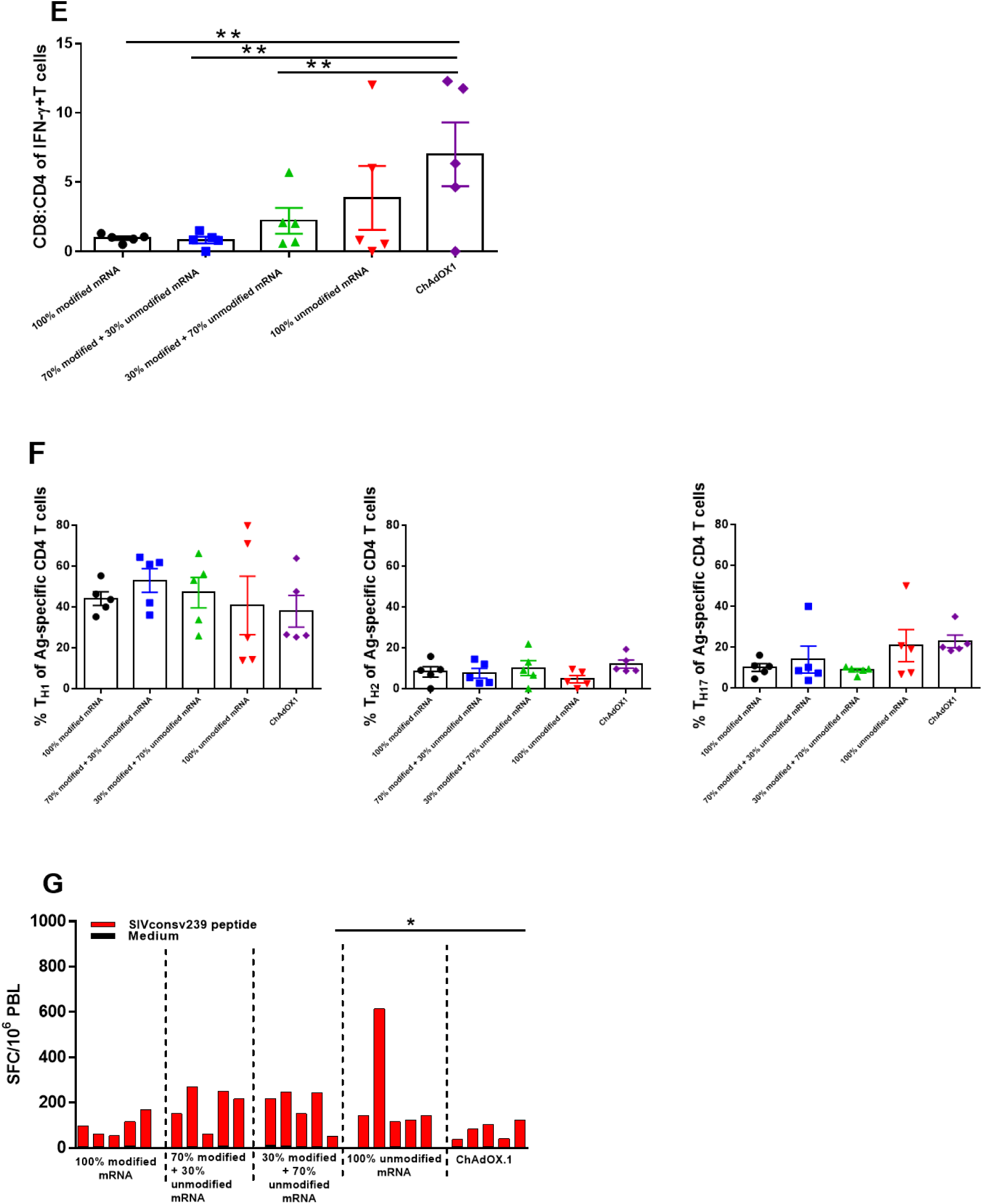
Immunization of rhesus monkeys with 100% unmodified mRNA expressing SIVconsv239 immunogen elicits higher T cell responses than 100% modified mRNA. Groups of monkeys (n=5/group) were immunized with mRNA-SIVconsv239 or ChAdOx1-SIVconsv239 immunogens. (**A**) CD8^+^ T cell responses specific for the immunodominant SIV Gag p11C epitope measured by tetramer staining of whole blood from the *Mamu-A*01*+ monkeys in each vaccination group at weeks 10 (blue bars), 12 (red bars) and 16 (green bars) of the study. The percent CD3^+^ CD8^+^ T cells that bound the *Mamu-A*01*/peptide-tetramer complexes in individual *Mamu-A*01*^+^ animals are shown. (**B**) PBMC IFN-γ ELISpot responses at study week 12, 4 weeks post final mRNA and 8 weeks post-ChAdOx1 immunizations. Freshly isolated PBMC were assessed for IFN-γ ELISpot responses after *in vitro* stimulation with a SIVconsv239 peptide pool spanning the entire immunogen sequence. Red bars represent antigen-specific and black bars represent the background spot forming cell (SFC) responses per million PBMC from individual monkeys in each group. **(C-D)** Cytokine secretion profiles of CD4^+^ **(C)** and CD8^+^ **(D)** T lymphocytes of the mRNA and ChAdOx1.SIVconsv239-vaccinated monkeys. Percent IFN-γ (top panels), IL-2 (middle panels) and TNF-α (bottom panels) positive CD4^+^ and CD8^+^ T lymphocytes from individual monkeys in each group following stimulation with a SIVconsv239 peptide pool are shown on study week 12, 4 weeks post final mRNA or ChAdOx1.SIVconsv239 immunization. **(E)** CD8:CD4 ratio within antigen-specific IFN-γ+ T cells in the five groups of immunized monkeys on study week 12**. (F)** Antigen-specific CD4^+^ T cells (defined on the basis of activation marker upregulation in response to SIVconsv239 peptide stimulation) were analyzed to determine their T_H1_/T_H2_/T_H17_ phenotypes. Percentages of antigen-specific T_H1_, T_H2_, and T_H17_ CD4^+^ T cells at study week 10 are shown in five groups of immunized monkeys. In panels C-F, symbols represent data from individual animals and mean ± SEM per group are indicated (**p<0.01; one-way ANOVA with Bonferroni correction). **(G)** PBMC IFN-γ ELISpot responses 26 weeks post final mRNA immunizations at study week 34. Freshly isolated PBMC were assessed for IFN-γ ELISpot responses after *in vitro* exposure to a SIVconsv239 peptide pool spanning the entire immunogen. Red bars represent antigen-specific and black bars represent the background SFC responses per million PBMC from individual monkeys in each group (*p< 0.05 Kruskal-Wallis test).

Median antigen-specific IFN-γ ELISpot responses analyzed at study week 12 also trended to be higher in the 100% unmodified mRNA group 4 (862 SFC/million PBMC) than in the 100% modified mRNA group 1 (397 SFC/million PBMC) (p= 0.121; Kruskal-Wallis test). Groups 4 and 5 had comparable spot forming cell (SFC) responses (p=0.22; Kruskal-Wallis test) (**Fig. 1B**). ICS assays performed at study week 12 showed significantly higher frequencies of IFN-γ-producing CD4^+^ T cells in group 4 than in group 1 and group 5 (Bonferroni-adjusted p <0.05; one-way ANOVA) (**Fig. 1C**, top panel), with monkeys in groups 2 and 3, immunized with mixtures of modified and unmodified nucleoside containing mRNA, showing IFN-γ-producing CD4 responses of intermediate magnitude. IL-2- and TNF-α-producing CD4^+^ T cell responses also trended to be higher in group 4 compared to group 1 (p=0.25; Kruskal-Wallis test) and 5 monkeys (p=0.125; Kruskal-Wallis test). No IL-4 or IL-10-producing T cell responses were detected (**Fig. S4**). Similarly, IFN-γ-producing CD8^+^ T cell frequencies in group 4 and group 3 trended to be higher than those in groups 1 and 2 (Bonferroni-adjusted p>0.05; one-way ANOVA). ChAdOx1-immunized monkeys also had high frequencies of cytokine-producing CD8^+^ T cells (**Fig. 1D**). However, frequencies of cytokine-producing CD4^+^ T cells in the ChAdOx1 group were relatively low compared to those in the unmodified mRNA-immunized groups, and analyses of the ratio of IFN-γ-secreting CD8^+^ to CD4^+^ T cells in each group at study week 12 revealed that this was significantly higher in ChAdOx1 immunized monkeys than in the group 1, 2 and 3 monkeys (Bonferroni-adjusted p>0.05; one-way ANOVA) immunized with mixtures of unmodified and modified nucleoside containing mRNA **(Fig. 1E).** In the mRNA-immunized groups, addition of increasing percentages of unmodified nucleoside containing mRNA increased the ratio of CD8^+^ to CD4^+^ T cells, with 100% unmodified mRNA inducing the most highly CD8^+^ T cell-biased T cell response.

We also analyzed the phenotypes of the antigen-specific CD4^+^ T cells by evaluating expression of chemokine receptors on CD4^+^ T cells that upregulated activation markers following *in vitro* stimulation with SIVconsv239 peptides (gating strategies for ICS and AIM assays are shown in **Fig. S3A-B**). As shown in **Fig. 1F**, in all five groups of monkeys, antigen-specific CD4^+^ T cells had a predominantly T_H1_ phenotype, with a relatively low proportion of T_H2_-phenotype cells. Monkeys immunized with either 100% unmodified mRNA or ChAdOx1 showed a trend to have a higher frequency of CD4^+^ T cells with T_H17_ phenotype compared to the other groups (p=0.69; Kruskal-Wallis test). Notably, IFN-γ assays performed 26 weeks after the last mRNA immunizations showed that unmodified mRNA immunizations elicited more durable antigen-specific T cell responses than two serial immunizations with ChAdOx1 (p <0.05; Kruskal-Wallis test) (**Fig. 1G**).

### Addition of IL-12-encoding mRNA to immunizations enhanced antigen-specific CD8^+^ T cell responses

Adding mRNA encoding IL-12 as an adjuvant has been reported to enhance CD8^+^ T cell function in different disease models including SARS-CoV-2 (*44, 46*). To augment the CD8^+^ T cells elicited by prior mRNA and ChAdOx1 immunizations, we added two booster SIVconsv239 mRNA-LNP immunizations that were delivered with 100% modified mRNA-LNP expressing a macaque IL-12p35-GS-p40 chimera (*44*) on weeks 41 and 58 of the study (**Table S2**). Monkeys in groups 1-4 received the same percentage of modified or unmodified nucleoside containing mRNA-SIVconsv239 used for the priming immunizations, in addition to 50 μg of modified mRNA-IL-12 subcutaneously. Monkeys primed with ChAdOx1 received 50 μg of 100% modified mRNA-SIVconsv239 with 50 μg of modified mRNA-IL-12. Longitudinal SIV-p11C tetramer-reactive CD8^+^ T cell responses in whole blood of the monkeys are shown in **Fig. 2A**. Time points where IL-12 adjuvant was added are shown on the x-axis with red arrows. Monkeys that were primed with 30%, 70% or 100% unmodified mRNA or ChAdOx1 showed a 1.5 to 11-fold increase in SIV Gag dominant epitope-specific CD8^+^ T cell responses relative to those present on study week 12. These responses were durable in 100% unmodified mRNA and ChAdOx1 primed groups and were still elevated relative to pre-boost frequencies at study week 78, which was 20 weeks after the final mRNA-IL-12 immunization **(Fig. 2A**). At study week 60, antigen-specific IFN-γ ELISpot responses in groups 3, 4 and 5 were significantly higher than those in group 1 (Bonferroni-adjusted p <0.05, one-way ANOVA), with particularly strong responses being observed in group 5 monkeys, which were primed with ChAdOx1 and given a heterologous vector boost with 100% modified mRNA encoding the SIVconsv239 immunogen, together with IL-12-encoding mRNA (**Fig. 2B**). Monkeys in group 5 also had the highest increase (12-120-fold) in magnitude of IFN-γ ELISpot responses following two IL-12 adjuvanted immunizations (**Fig. S5**). ICS assays performed on PBMCs of all monkeys at study week 59, which was one week post-2^nd^ SIVconsv239-mRNA + IL-12-mRNA booster immunization, showed that IFN-γ-producing CD4^+^ T cell frequencies in group 4 monkeys primed and boosted with 100% unmodified mRNA were significantly higher (p <0.05; Kruskal-Wallis with Dunn’s post-test) than those in group 1 animals immunized with 100% modified mRNA **(Fig. 2C**). A trend for higher responses relative to group 1 was also observed in group 3 monkeys immunized with 70% unmodified mRNA (p= 0.155; Kruskal-Wallis test) **(Fig 2C)**. IFN-γ, IL-2 and TNF-α-producing CD8^+^ T cell frequencies were also significantly higher in group 4 than group 1 (p= 0.015, 0.01 and 0.0076, respectively, Kruskal-Wallis with Dunn’s post-test), with group 3 showing a similar trend for higher responses relative to those of group 1 **(Fig 2D).** Group 5 monkeys primed with ChAdOx1 and boosted with 100% modified mRNA expressing the SIVconsv239 immunogen also had significantly higher frequencies of IFN-γ and TNF-α- producing CD8^+^ T cells than group 1 monkeys that were both primed and boosted with 100% modified mRNA-SIVconsv239 (p=0.008; Kruskal-Wallis with Dunn’s post-test) (**Fig. 2D**).

**Fig. 2.**
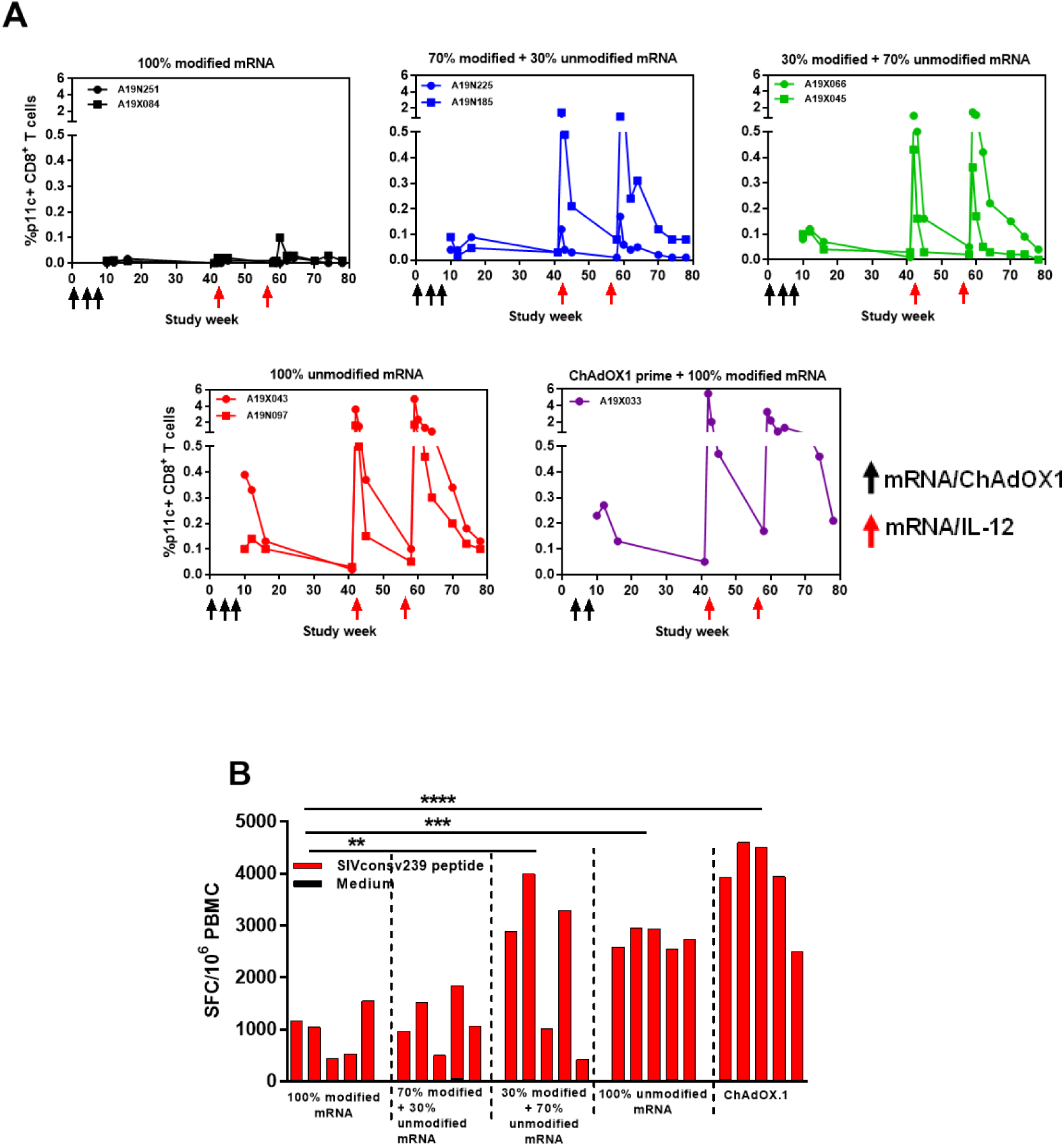
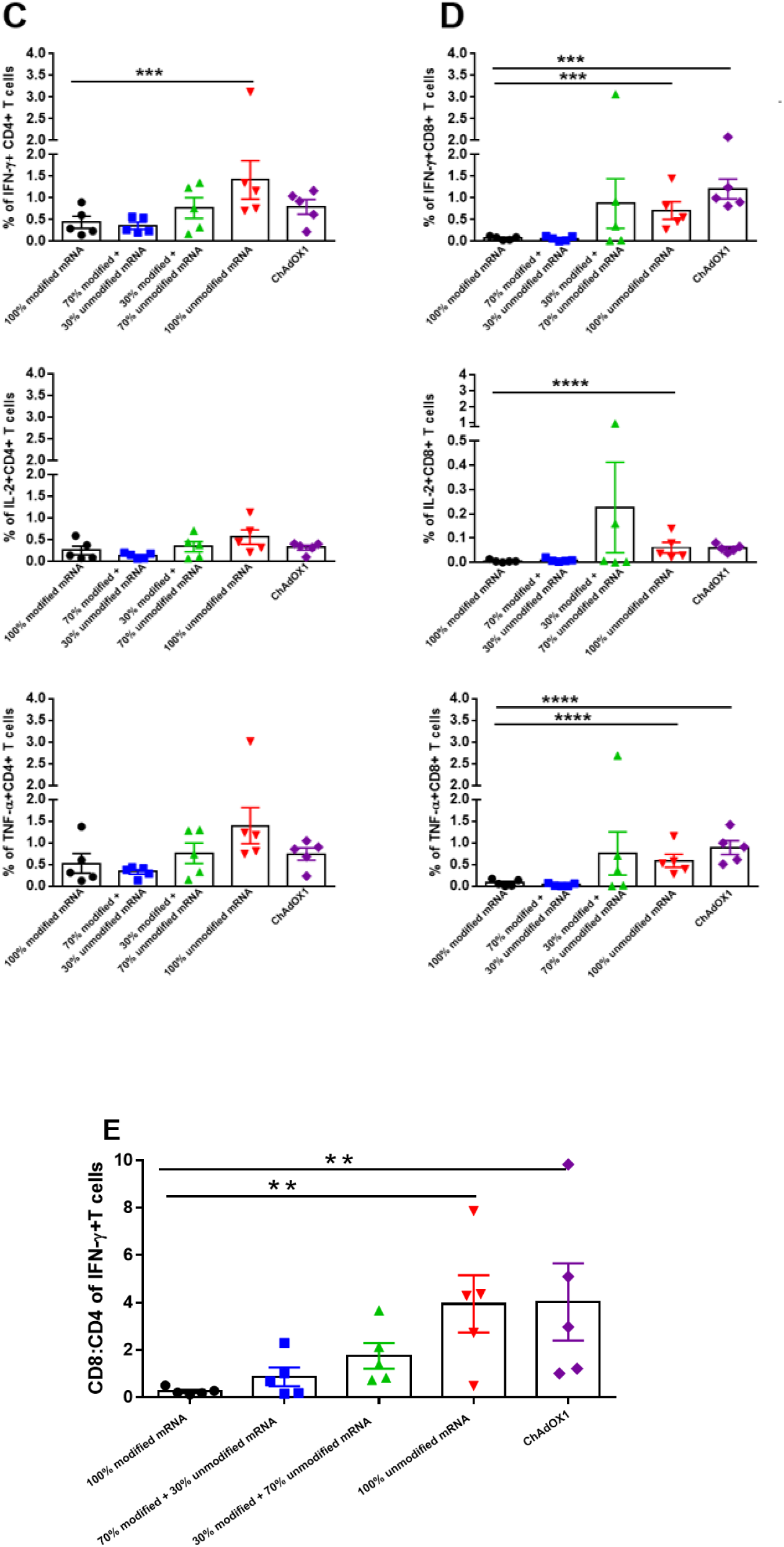
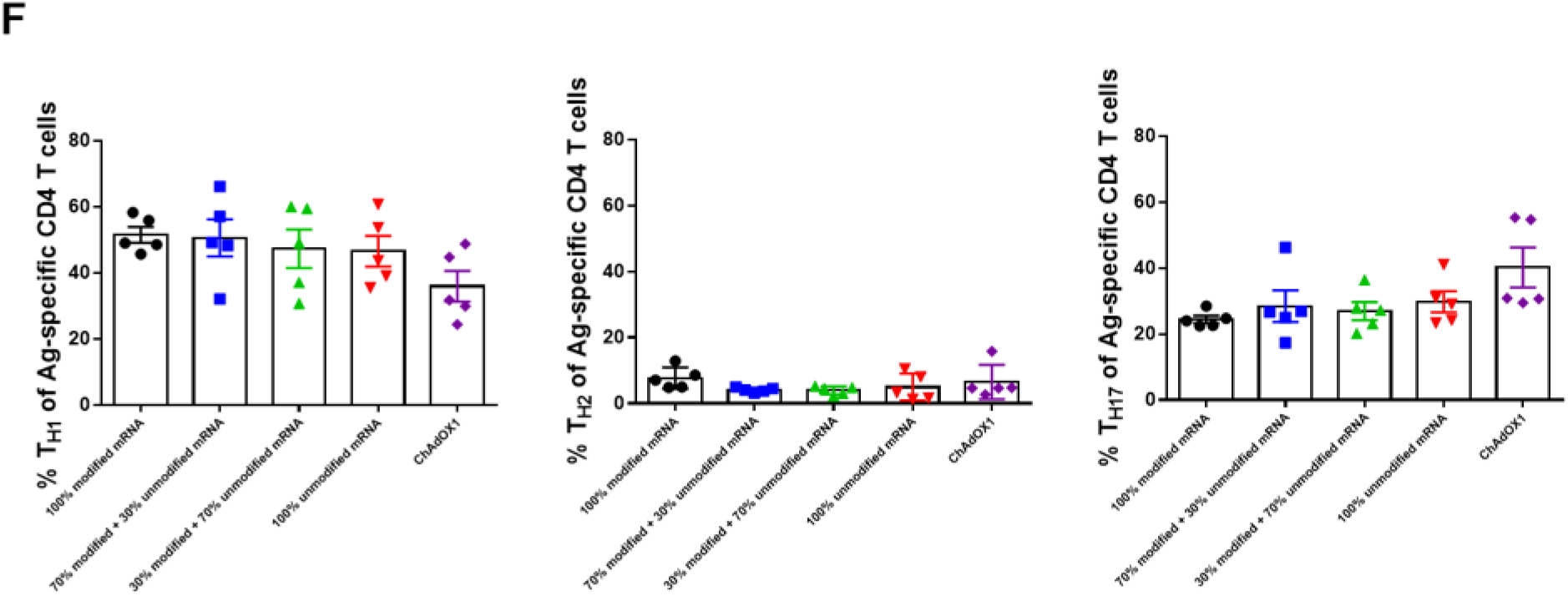
Booster SIVconsv239-mRNA immunizations administered with IL-12-mRNA induce very high magnitude antigen-specific CD8^+^ T cell responses in both SIVconsv239-mRNA and ChAdOx1.SIVconsv239-primed monkeys. **(A)** CD8^+^ T cell responses specific for the immunodominant SIV Gag p11C epitope measured by tetramer staining of whole blood from the *Mamu-A*01*^+^ monkeys in each vaccination group through the entire course of the study. Black arrows indicate priming immunizations, and red arrows indicate boosting immunizations administered with modified IL-12 mRNA-encoding LNPs. The percent CD3^+^ CD8^+^ T cells that bound the *Mamu-A*01*/peptide-tetramer complexes in individual *Mamu-A*01*+ animals are shown. **(B)** PBMC IFN-γ ELISpot responses at study week 60, 2 weeks after final mRNA-IL-12 immunization. Freshly isolated PBMC were assessed for IFN-γ ELISpot responses after *in vitro* stimulation with a SIVconsv239 peptide pool spanning the entire immunogen sequence. Red bars represent antigen-specific and black bars represent the background SFC responses per million PBMC from individual monkeys in each group (**p<0.01; ***p<0.001; ****p<0.0001; one-way ANOVA with Bonferroni correction). **(C-D)** Cytokine secretion profiles of CD4^+^ **(C)**, CD8^+^ **(D)** T lymphocytes of the mRNA and ChAdOx1.SIVconsv239-vaccinated monkeys. Percent IFN-γ (top panels), IL-2 (middle panels) and TNF-α (bottom panels) positive CD4^+^ and CD8^+^ T lymphocytes from individual monkeys in each group following stimulation with a SIVconsv239 peptide pool on week 59 of the study are shown and the group mean ± SEM are indicated (***p<0.001; ****p<0.0001; Kruskal-Wallis with Dunn’s post-test). **(E)** CD8:CD4 within antigen-specific IFN-γ^+^ T cells in five groups of immunized monkeys. Mean ± SEM per group are shown (**p<0.05; one-way ANOVA with Bonferroni correction). (**F**) Percentages of antigen-specific T_H1_, T_H2_, and T_H17_ CD4^+^ T cells at study week 59 are shown in five groups of immunized monkeys and the group mean ± SEM are indicated.

These CD8^+^ T cells also exhibited cytotoxic potential, as shown by granzyme B staining and analysis of degranulation capacity (CD107a expression) (**Fig. S6**). We also determined the ratio of IFN-γ-secreting CD8^+^ and CD4^+^ T cells post-boosting with concurrent IL-12-mRNA immunization. Both 100% unmodified mRNA and ChAdOx1-primed monkeys had a significantly higher ratio of IFN-γ secreting CD8^+^ to CD4^+^ T cells than the 100% modified mRNA primed group **(Fig. 2E)** (Bonferroni-adjusted p <0.05, one-way ANOVA). In addition, we analyzed the phenotypes of the antigen-specific CD4^+^ T cells based on expression of chemokine receptors (gating strategy in **Fig. S3**). As shown in **Fig. 2F**, in all five groups of monkeys, antigen-specific CD4^+^ T cells had a predominantly T_H1_ rather than a T_H2_ phenotype, with some T_H17_-phenotype cells also being present. The pattern of T_H1_/T_H2_/T_H17_ phenotypes was not appreciably altered following mRNA-IL-12 administration.

We collected inguinal lymph node biopsies from all five groups of monkeys one-week post-2^nd^ mRNA-IL-12 immunization on week 59 and performed ICS assays on lymph node lymphocytes. As shown in **Fig. S7**, cytokine secreting CD4^+^ T cells (top panels) trended to be highest in groups primed with 100% or 70% unmodified mRNA (p=0.15; Kruskal-Wallis with Dunn’s post-test), whereas cytokine^+^ CD8^+^ T cells (bottom panels) trended to be higher in ChAdOx1-primed monkeys (p=0.14; Kruskal-Wallis with Dunn’s post-test).

### Circulating CD4^+^ T follicular helper cell frequencies correlated with SIV Gag-specific binding antibody responses

We also analyzed frequencies of circulating T_FH_ -phenotype CD4^+^ T cells, (cT_FH_, defined as PD1^+^CXCR5^+^CD4^+^ T cells) in peripheral blood on week 10 and week 59 of the study (**Fig. 3A- B**). At week 10, monkeys immunized with 100, 70 or 30% modified mRNA showed a trend for higher percentages of cT_FH_ cells compared to the 100% unmodified mRNA and ChAdOx1 primed groups (p=0.59; Kruskal-Wallis with Dunn’s post-test). Importantly, at week 59, cT_FH_ frequencies in group 1 monkeys (primed and boosted with 100% modified mRNA), were significantly higher than those in groups 4 (primed and boosted with 100% modified mRNA) and 5 (primed with ChAdOx1, boosted with 100% modified mRNA) [Bonferroni-adjusted p=0.01; one-way ANOVA].

**Fig. 3.**
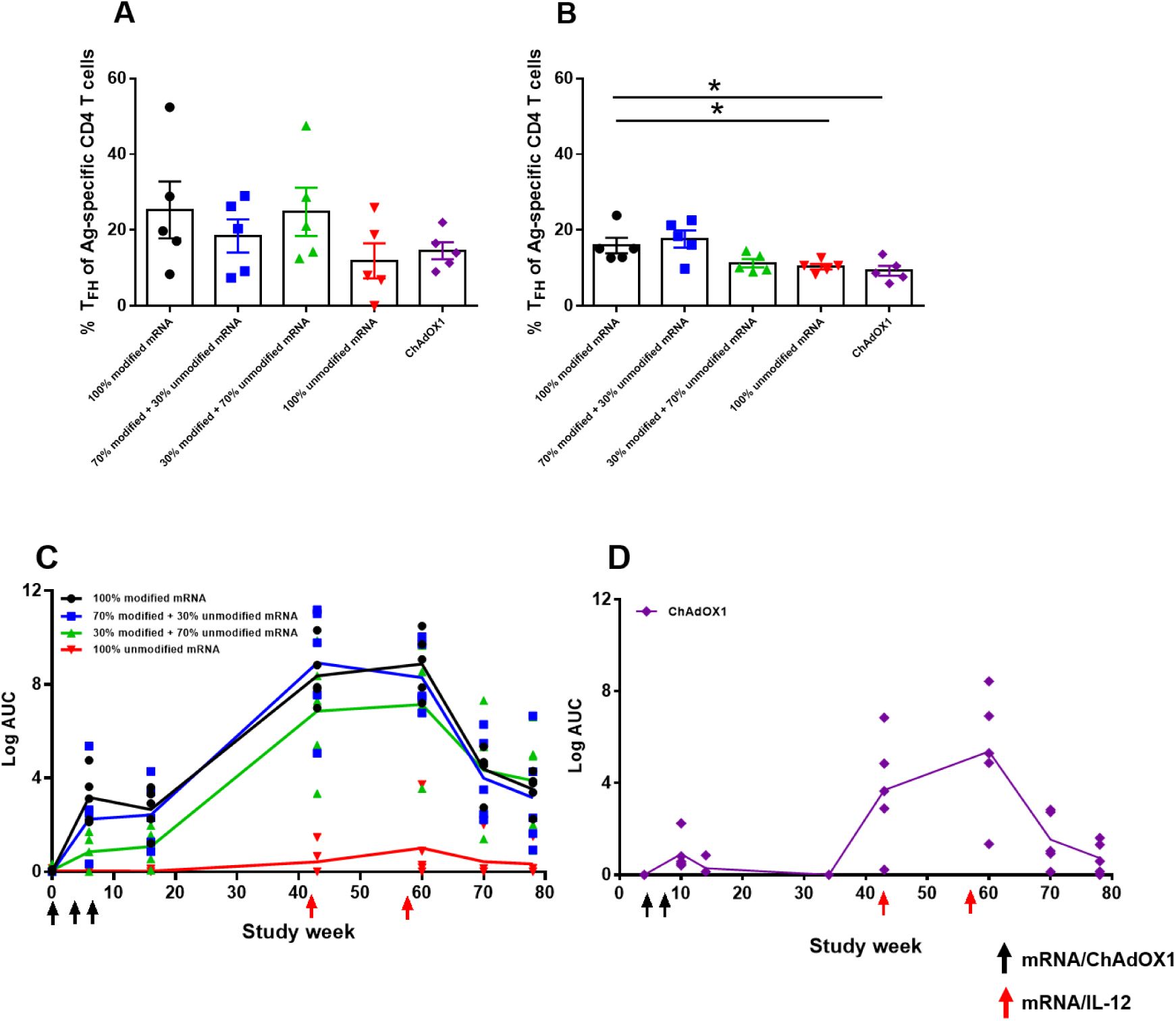
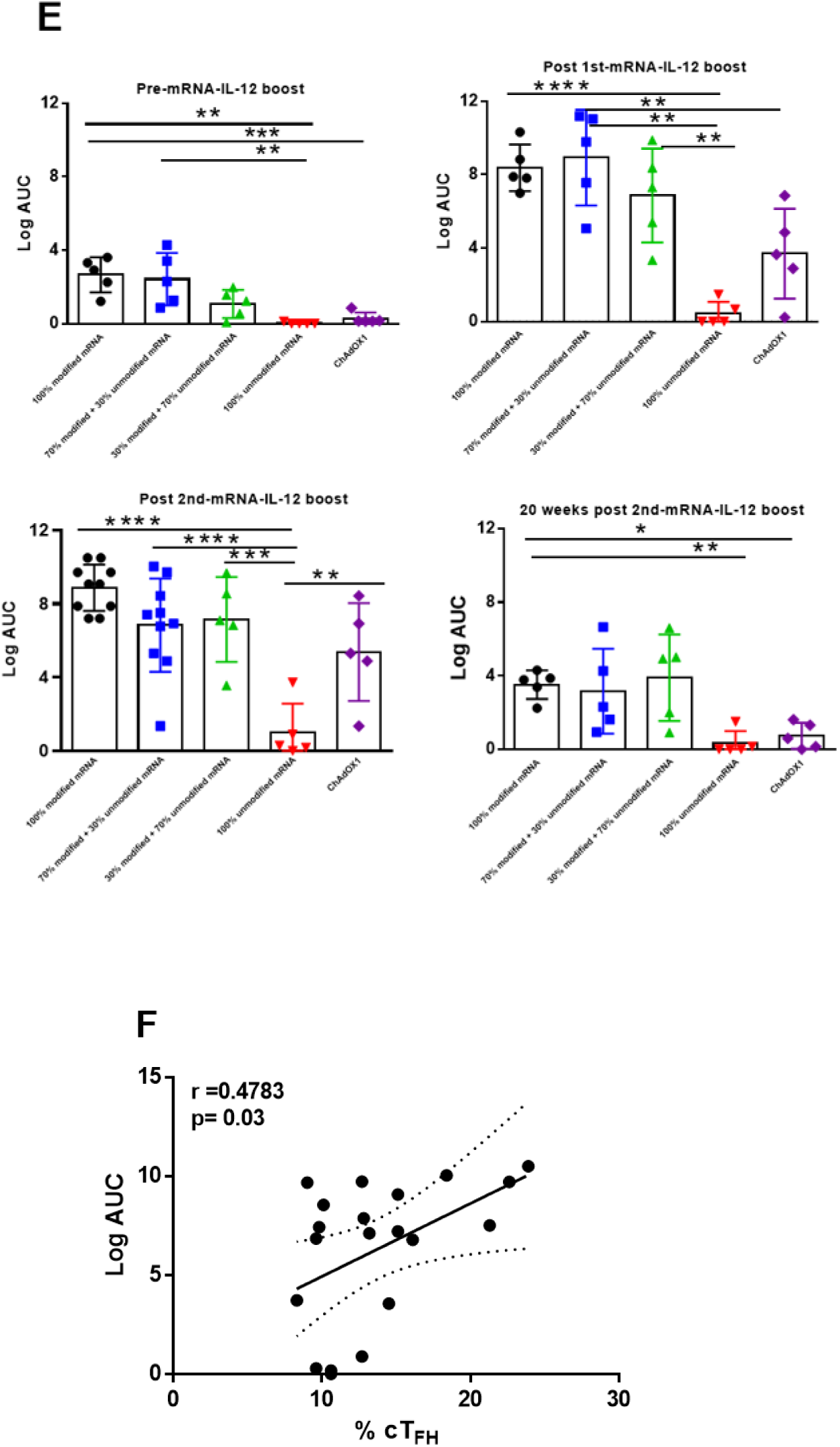
Immunization with 100% modified mRNA elicits higher CD4^+^ T follicular helper cell (T_FH_) responses and longitudinal SIV Gag-specific binding antibody responses than 100% modified mRNA or ChAdOx1-vectored immunogen. Percentages of antigen-specific CD4^+^ T_FH_ CD4^+^ T cells at study weeks 10 **(A)** and 59 **(B)** are shown in the five groups of immunized monkeys. Mean ± SEM per group are indicated (* p<0.05; one-way ANOVA with Bonferroni correction). **(C-D)** Plasma from individual monkeys was tested for the presence of Abs binding to SIV Gag protein by ELISA. Log area under the curve (AUC) of the binding titers in each individual monkey in the four groups primed three times with different ratios of mRNA (**C**) and the group primed twice with ChAdOx1 (**D**) are shown (symbols). Each line represents the median binding titer of the five monkeys in an immunization group (groups are shown in different colors). Immunization time points are shown on the x-axis. **(E)** Log AUC of the Gag-specific Ab binding titers per monkey in each group are shown for pre-mRNA-IL-12 boost (study week 16) (upper left), two weeks post-1st mRNA-IL-12 (upper right) or post-2^nd^ mRNA-IL-12 boost (lower left), and 20 weeks post-2^nd^ mRNA-IL-12 boost (lower right) immunizations. Mean + SD values for each group are shown (*p <0.05; **p<0.01; ***p<0.001; ****p<0.0001; Kruskal-Wallis with Dunn’s post-test). **(F)** Correlation between circulating T follicular helper cells (% cT_FH_) and antibody response at study week 60 (Log AUC). The solid line shows the least-squares linear regression and the area denoted with dotted lines represents the 95% confidence interval.

We next measured SIV Gag-specific binding antibody (Ab) responses longitudinally by ELISA in these five groups of immunized monkeys. Binding Ab titers (log_10_ area under the curve (AUC) values) of individual monkeys and group medians of the groups primed with mRNA are shown in **Fig. 3C**. As the ChAdOx1 immunizations started four weeks after groups 1-4, group 5 responses are shown in **Fig 3D**. Monkeys in groups 1 and 2, which were primed and boosted with 100% and 70% modified mRNA, respectively, had the highest plasma binding Ab responses throughout the study. Notably, antibody responses in these groups increased 2-9-fold after the boosting immunizations, which were given with IL-12-encoding mRNA (**Fig. 3C)**. In group 4 monkeys, immunized with 100% unmodified mRNA, antibody responses were scarcely detected after priming, and only a low-level response was observed after the IL-12 adjuvanted boosts. Responses in group 1 and 2 monkeys (primed and boosted with 100% and 70% modified mRNA, respectively) were significantly higher than those in group 4 monkeys (primed and boosted with 100% unmodified mRNA) at both pre-and post-boost timepoints (p<0.01 for groups 1 vs 4 and group 2 vs 4 responses at both pre- and post-boost time points; Kruskal-Wallis with Dunn’s post-test) (**Fig. 3E**). Post-priming, group 3 animals (immunized with 30% modified mRNA) had an antibody response of intermediate magnitude; and this was increased by 2-10 fold after boosting plus IL-12 mRNA (**Fig. 3C**), bringing the magnitude of responses comparable to groups 1 (100% modified mRNA) and 2 (70% modified mRNA). Two weeks after the 1^st^ and 2^nd^ mRNA boosts responses in not only groups 1 and 2 but also group 3 animals were significantly higher than those in group 4 (group 3 vs group 4 p<0.01 and p<0.001 after the 1^st^ and 2^nd^ boosts, respectively; Kruskal-Wallis with Dunn’s post-test) (**Fig. 3E**). This finding was also maintained as a trend at late post-boost timepoints (p<0.001; Kruskal-Wallis with Dunn’s post-test). Priming with ChAdOx1 elicited very low binding antibody titers, but although responses increased 22-50-fold after boosting with 100% modified mRNA with IL-12-encoding mRNA, they remained significantly lower than those in group 1 at study week 60 (p<0.05; Kruskal-Wallis test) (**Fig. 3E)**.

We also addressed whether the frequencies of cT_FH_ -phenotype cells induced following priming with modified and/or unmodified mRNA correlated with subsequent Gag-specific binding Ab titers. A weak positive correlation was observed between the week 10 cT_FH_ frequencies in animals in groups 1-4 and their Gag-specific binding Ab titers at week 60 of the study (r=0.26; Pearson correlation). However, significant positive correlations were observed between the week 59 cT_FH_ frequencies in animals in groups 1-4 and their Gag-specific binding Ab titers at study week 60 (r=0.48, p=0.031; Pearson correlation) (**Fig. 3F**).

### Peripheral and lymph node cellular immune responses elicited by mRNA-SIVconsv239 prime and mRNA-IL-12 boost are durable

Cellular immune responses were followed for 20 weeks post final mRNA-IL-12 boost (study week 78) to determine the durability of the vaccine-elicited responses. At this timepoint, groups 1 and 4 monkeys were necropsied to collect peripheral blood, axillary and inguinal lymph nodes and terminal ileum. As shown in **Fig. 2A**, *MamuA*01*-p11C tetramer-reactive CD8^+^ T cells were still detectable in the peripheral blood in group 4 and 5 monkeys at study week 78.

Although the magnitude of peripheral blood SIVconsv239-specific IFN-γ-producing T cell responses as assessed by ELISpot assay had contracted relative to study week 16 values, robust responses were still present at study week 78 (**Fig. 4A**). ELISpot responses and IFN-γ-secreting CD4^+^ (**Fig. 4B**) and CD8^+^ (**Fig. 4C**) T cell frequencies as measured by ICS in groups 3, 4 and 5 trended to be higher than those in groups 1 and 2, with IFN-γ ELISpot responses in group 5 monkeys being significantly higher than those in group 1 (Bonferroni adjusted p=0.009, one-way Anova) (**Fig. 4A**). Similarly, analysis of the frequency of SIVconsv239-specific cytokine-producing T cells in the axillary and inguinal lymph nodes revealed the presence of significantly higher frequencies of IFN-γ-producing CD4^+^ and CD8^+^ T cells in the axillary lymph nodes in group 4 monkeys immunized with 100% unmodified mRNA (**Fig. 4D**) than in the group 1 animals immunized with 100% modified mRNA (p<0.05; Mann-Whitney U test). These results were paralleled in the inguinal lymph nodes (**Fig. 4E**), with lymph node IL-2- and TNF-α-producing CD4^+^ and CD8^+^ T cell frequencies showing the same trends.

**Fig. 4.**
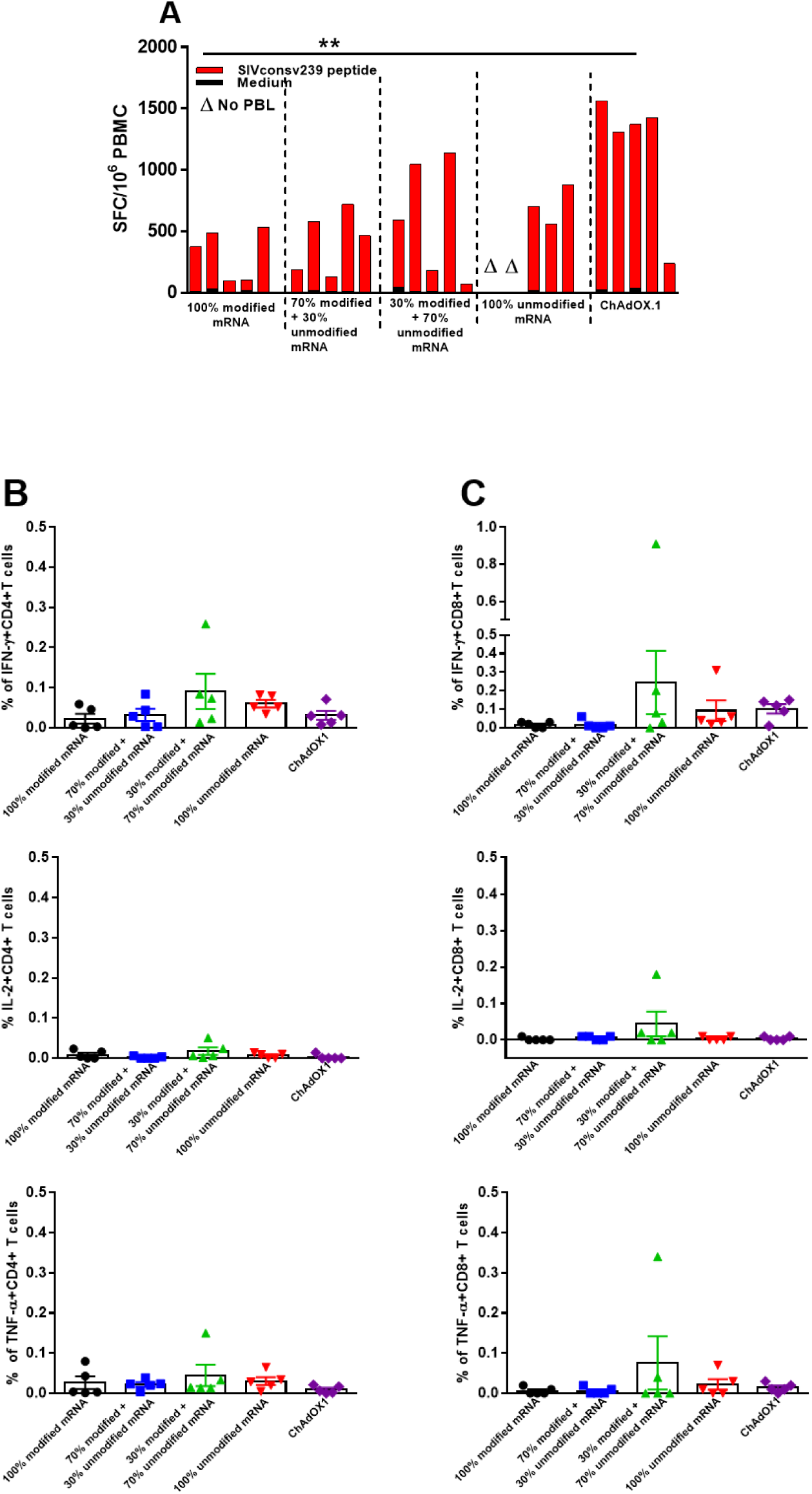
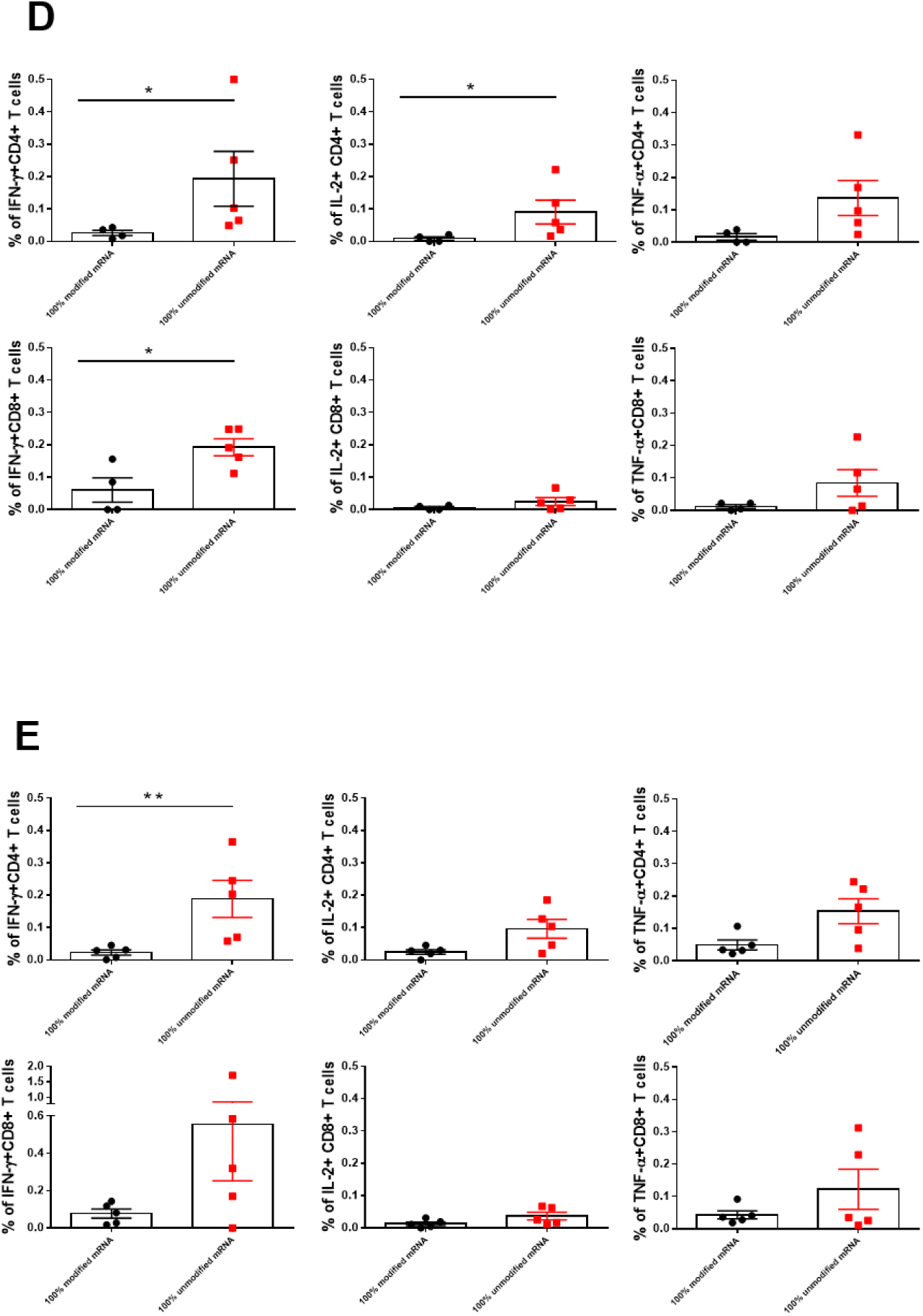
Durability of the cellular immune responses in groups of rhesus monkeys after boosting with different ratios of modified and unmodified mRNA, co-administered with IL-12-mRNA. **(A)** PBMC IFN-γ ELISpot responses 20 weeks after the second booster immunization, given with IL-12-mRNA. Freshly isolated PBMC were assessed for IFN-γ ELISpot responses after *in vitro* stimulation with a SIVconsv239 peptide pool spanning the SIVconsv239 immunogen sequence. Red bars represent antigen-specific and black bars represent the background SFC responses per million PBMC from individual monkeys in each group. Data are shown for only 3 of the 5 monkeys in the 100% unmodified mRNA-immunized group as two animals had insufficient cells for analysis (Δ). (**p<0.01; one-way ANOVA with Bonferroni correction) (**B-C**) Cytokine secretion profiles of CD4^+^ (**B**) and CD8^+^ (**C**) T lymphocytes of the monkeys in each group. Percent IFN-γ (top panels), IL-2 (middle panels) and TNF-α (bottom panels) positive CD4^+^ and CD8^+^ T lymphocytes from individual monkeys in each group following stimulation with a SIVconsv239 peptide pool on week 78 of the study are shown. (**D-E**) Cytokine secretion profile of lymph node CD4^+^ and CD8^+^ T cells. Monkeys immunized with either 100% modified mRNA (group 1) or 100% unmodified mRNA (group 4) were necropsied at week 78 of the study. Lymphocytes isolated from axillary (**D**) and inguinal **(E)** lymph nodes were assessed for their cytokine secretion profiles. Secretion of cytokines by CD4^+^ and CD8^+^ lymph node T cells are shown. In panels B-E, symbols represent data from individual animals, and bars show mean ± SEM per group. In panels D and E, *p<0.05; **p<0.01; Mann-Whitney U test.

## DISCUSSION

Here, we show in rhesus macaques that serial priming and boosting with unmodified mRNA in LNPs elicited higher magnitude CD8^+^ T cell responses than modified mRNA as analyzed by MHC class I-peptide tetramer staining of whole blood, and antigen-specific IFN-γ ELISpot assays and ICS assays. Antigen-specific CD8^+^ T cell responses were significantly higher in the group of macaques serially primed and boosted with 100% unmodified mRNA than in the group serially primed and boosted with 100% modified mRNA.

A 30% modified/70% unmodified nucleoside containing mRNA optimized both CD8^+^ T cell, CD4^+^ T_FH_ and antibody responses. Boosting immunizations were co-administered with mRNA encoding IL-12 and drove substantial expansion of both T cell and antibody responses.

A previous study in mice comparing responses elicited by modified and unmodified mRNA-LNP vaccines encoding ovalbumin or tumor neoantigens also suggested induction of superior CD8^+^ T cell responses and greater therapeutic efficacy by unmodified mRNA vaccines (*36*), and a recent study in mice evaluating the immunogenicity of modified and unmodified mRNA-LNP vaccines encoding Mycobacterium tuberculosis antigens similarly reported higher levels of polyfunctional and memory CD8^+^ cells in the unmodified mRNA-LNP vaccinated group (*47*). In the former study, the superior CD8^+^ T cell responses and therapeutic efficacy of unmodified mRNA vaccines were shown to depend on their induction of high levels of type I IFNs (*36*), innate cytokines that support induction and maintenance of CD8^+^ T cell responses via effects on both dendritic cells (DCs) and T cells (*48*) but reduce mRNA translation and humoral responses (*49–51*). In our study, we demonstrate more potent induction of antigen-specific CD8^+^ T cell responses by 100% unmodified than 100% modified mRNA-LNP vaccines in non-human primates, and, as discussed further below, define the proportions of modified and unmodified nucleoside containing mRNA-LNP that induce both high levels of CD8^+^ T cells and high-magnitude serum antibody responses.

Our study included a comparator group of rhesus macaques primed with two immunizations of ChAdOx1.SIVconsv239, an adenoviral vector previously shown to induce strong CD8^+^ T cell responses in both pre-clinical and clinical settings (*41, 52–54*). Priming with three immunizations of 100% unmodified SIVconsv239-mRNA elicited antigen-specific CD8^+^ T cell responses of a comparable magnitude to this viral vector, and these were accompanied by a stronger antigen-specific IFN-γ-producing CD4^+^ T cell response. These CD8^+^ T cell responses were further expanded after boosting, paralleling the expansions observed in the ChAdOx1.SIVconsv239-primed monkeys, which were boosted with 100% modified SIVconsv239-mRNA. This heterologous boost not only enhanced antigen-specific CD8^+^ T cell responses but also increased the accompanying CD4^+^ T cell response, making the antigen-specific IFN-γ-producing CD8:CD4 T cell ratio more comparable to that in animals both primed and boosted with 100% unmodified SIVconsv239-mRNA. Given the importance of CD4^+^ T cells in the establishment and maintenance of CD8^+^ T cell memory (*55*), the induction of strong antigen-specific CD4^+^ T cell responses likely contributed to the durability of the CD8^+^ T cell responses in both the heterologous prime-boosted group and monkeys serially primed and boosting with 100% unmodified SIVconsv239-mRNA.

Although serial priming and boosting with 100% unmodified mRNA elicited significantly higher antigen-specific CD8^+^ and CD4^+^ T cell responses than priming and boosting with 100% N1-methylpseudouridine modified mRNA, Gag-specific antibody responses were scarcely detected after priming with 100% unmodified mRNA and only a low-level response was observed after the IL-12 co-administered boost. However, there was a greater CD4:CD8 cell bias within antigen-specific IFNγ-producing T cells and a significantly higher frequency of cT_FH_ cells within the antigen-specific CD4^+^ T cell population in the group primed and boosted with 100% modified mRNA, and significantly higher antibody responses were observed at both pre- and post-boost timepoints (which were paralleled in the group immunized with 70% modified mRNA). Other comparative studies in pre-clinical models have similarly found mRNA immunogens containing increasing percentages of nucleoside-modified mRNA to elicit higher-titer antibody responses, associated with decreased innate sensing and enhanced mRNA translation (*37, 49–51*). The influence of levels of innate activation and antigen synthesis on the magnitude of the antibody response elicited by modified versus unmodified mRNA-LNP vaccines is highlighted by two studies in macaques, one of which showed that repeated immunization with high doses of a modified mRNA-LNP vaccine that stimulated similarly-robust (although qualitatively-different) innate activation to repeated immunization with a lower dose of an unmodified Gag mRNA-LNP vaccine elicited Gag-specific antibody responses of only comparable magnitude to those induced in the unmodified Gag mRNA-LNP vaccinated group (*56*). The second study showed the optimization of the non-coding regions of the CVnCoV (CureVac) unmodified mRNA-LNP SARS-CoV-2 vaccine to enhance translation enabled induction of substantially higher antibody responses in macaques, rendering this unmodified mRNA-LNP vaccine similarly immunogenic to the BNT162b2 (Pfizer) modified mRNA SARS-CoV-2 vaccine encapsulated in the same LNPs (*57*).

Boosting immunizations of SIVconsv239-mRNA-LNP vaccines were co-administered with IL-12-mRNA-LNPs. At the time of its discovery, IL-12 was identified to be a product of Epstein–Barr virus-transformed human B cell lines that can activate NK cells and induce IFN-γ production and T cell proliferation (*58–60*). Subsequent reports demonstrated that IL-12 enhances IFN-γ production by T lymphocytes and NK cells (*61, 62*) and polarizes naive CD4^+^ T cells to become T_H1_ cells (*60, 61, 63*). IL-12 has also been shown to enhance CD8^+^ T cell proliferation and full activation, and to support survival of activated CD8^+^ T cells and memory formation (*64–68*). These T cell activating properties of IL-12 have led to its testing as a vaccine adjuvant (*69–71*). In the current study, we used mRNA-encoded IL-12 as a boost to enhance initially primed T cell responses to SIV immunogen, as systemic administration of IL-12 protein has been shown to be poorly tolerated (*72*). Our data show expansion of strong antigen-specific CD8^+^ and T_H1_-type CD4^+^ T cell responses with IFN-γ production following boosts with IL-12-mRNA, and no induction of T_H2_ cytokines. IL-12 and the IFN-γ it induces also act on B cells to promote proliferation and plasmablast differentiation (*73*), in line with serum antibody responses in the animals primed with 100%, 70% or 30% modified nucleoside containing mRNA-LNPs were substantially enhanced after boosting, and a low-magnitude antibody response was also detected in the 100% unmodified mRNA-primed group. Similarly, augmentation of humoral as well as cellular responses to a SARS-CoV-2-mRNA-LNP vaccine by co-administration with IL-12-mRNA-LNPs has been observed in mice (*46*).

An important finding in this study is the co-induction of both robust CD8^+^ T cell and CD4^+^ T cell and antibody responses in the group of animals serially primed and boosted with the 30% modified mRNA/70% unmodified nucleoside containing mRNA mixture. Two weeks after the first and second booster immunizations (administered with IL-12-mRNA), IFN-γ-producing CD8^+^ and CD4^+^ T cell responses in the 30% modified mRNA/70% unmodified mRNA-immunized group were of a comparable magnitude to those in the group immunized with 100% unmodified mRNA. Likewise, a study by Sittplangkoon *et al*. (*36*) in a mouse tumor model system demonstrated that immunization with a mixture of 40% modified/60% unmodified mRNA-LNPs induced similarly-high frequencies of antigen-specific IFN-γ-producing CD8^+^ T cells to those in mice immunized with 100% unmodified mRNA-LNPs encoding the same antigen, with significantly higher percentages of granzyme B and IFN-γ/granzyme B-producing CD8+ T cells being observed in these groups compared to those in animals immunized with 70 or 100% modified nucleoside containing mRNA-LNPs (*36*). Notably, here we found in non-human primates that serum Gag-specific antibody responses were significantly higher in the 30% modified mRNA/70% unmodified nucleoside containing mRNA-immunized group and stabilized at levels similar to those in the group immunized with 100% modified mRNA by 20 weeks after the second mRNA boost. In the absence of an envelope immunogen in the current vaccine study, we were unable to assess elicitation of neutralizing antibody, but our findings suggest benefits to this admixture for vaccines where induction of both CD8^+^ T cell and humoral immunity is needed for optimal protection, such as SARS-CoV-2 (*30–33*) or HIV-1(*74–77*). In future studies, it would be important to address whether contralateral delivery of 100% unmodified mRNA in one limb and 100% unmodified mRNA in the other limb would elicit comparable or better T cell and humoral immune responses to a mixed-composition mRNA-LNP vaccine.

The SARS-CoV-2 pandemic established the mRNA-LNP vaccine platform as a powerful tool for infectious diseases prevention, and it has subsequently also been applied to cancer vaccine development (*34, 78–81*). The ease of manufacturing and relative lack of reactogenicity make mRNA-LNPs prime candidates for HIV vaccine development (*82*). An array of HIV-1 antibody-inducing mRNA-LNP vaccine candidates are being evaluated in pre-clinical and clinical trials (*83–87*).

Data from this non-human primate study will inform optimal formulation of mRNA in LNPs to generate potent HIV-1-specific CD8^+^ T cell responses to provide an important additional means of HIV-1 vaccine-mediated protection in humans.

## MATERIALS AND METHODS

### SIVconsv239-mRNA and IL-12-mRNA constructs

mRNA design and production. Plasmids encoding SIVconsv239 immunogens were designed **(Fig. S1A-B)** and cloned for use as a template for in vitro transcription (Genscript). A plasmid-encoded 101-nucleotide polyadenylation sequence was included in the in vitro transcription template. Plasmids were linearized using Not-I, Afl-II, or Eco-RI endonucleases (NEB) and purified as previously described. mRNA immunogens were in vitro transcribed using T7 RNA polymerase (Megascript) with co-transcriptional capping (CleanCap 3OMe, Trilink) and incorporation of N1-methylpseudouridine in place of uridine (Trilink) implementing methods as previously reported (*83, 88*). For each condition with different N1-methylpseudouridine percentages, the remainder of the T/U content comprised UTP. Contaminating double stranded byproducts were removed with cellulose purification as previously described (*89*) and innate immune sensing was assessed as interferon-alpha release following transfection of monocyte-derived dendritic cells (*49*). mRNA was evaluated for appropriate size using gel electrophoresis and stored at -20°C. mRNA construct expressing IL-12 was described in Aunins et al (*44*) **(Fig. S1D)**.

### mRNA-LNP synthesis

mRNA-LNPs were prepared in 4-component lipid nanoparticles using methods as previously described (*90, 91*). LNP formulations contain ionizable lipid (proprietary to Acuitas)/phosphatidylcholine/cholesterol/PEG-lipid. Lipid and LNP compositions are described in US patent US10,221,127.

### ChAdOx1.SIVconsv239

ChAdOx1.SIVconsv239 is a replication-deficient vaccine designed to express the SIVconsv239 immunogen. The vaccine vector, ChAdOx1, is derived from the group E simian adenovirus Y25, originally isolated from a chimpanzee, and was specifically engineered for use in vaccine development (*40, 92*). The SIVconsv239 immunogen comprises six conserved regions from the Gag and Pol proteins of the SIVmac239 strain. Due to the limited availability of simian virus sequences, identifying conserved regions shared across most SIV variants remains challenging. To overcome this, SIVconsv239 incorporates protein segments that are homologous to the conserved Gag and Pol regions utilized in the candidate human HIVconsvX vaccines (*39*) **(Fig S1A-C**).

### Nonhuman primate vaccine study design

Monkeys were housed at the New Iberia Research Center (New Iberia, LA) and were maintained in accordance with the *Guide for the Care and Use of Laboratory Animals* with the approval of the Institutional Animal Care and Use Committee of University of Louisiana at Lafayette and the National Institutes of Health. Twenty-five Indian-origin rhesus (*Macaca mulatta*) monkeys were distributed in five experimental groups (n=5 per group) and were immunized as described in **Tables S1 and S2**. Monkeys in group 1-4 were primed with either 50 μg of 100% 1-methylpseudouridine modified SIVconsv239-mRNA, 70% modified SIVconsv239-mRNA + 30% unmodified SIVconsv239-mRNA, 30% modified mRNA+ 70% unmodified mRNA or 100% unmodified SIVconsv239-mRNA at weeks 0, 4, and 8. Monkeys in group 5 were primed with 10^10^ v.p. of ChAdOx1.SIVconsv239 at weeks 0 and 4. On weeks 41 and 58 of the study, monkeys in Groups 1-4 received boost with the same mixtures of mRNA-SIVconsv239 immunogens as prime as well as 50 μg of mRNA expressing IL-12. Monkeys in group 5 were boosted with 50 μg of 100% modified mRNA-SIVconsv239 and 50 μg of IL-12-mRNA. Blood samples were collected at pre-immunizations and every 2-4 weeks post-immunization time points. Lymph node biopsies were collected at week 59 of the study. Monkeys in groups 1 and 4 were necropsied on week 78 of the study to collect blood, lymph nodes and terminal ileum samples.

### IFN–γ ELISpot assay

Multiscreen ninety-six well plates were coated overnight with 100 μl per well of 5 μg/ml anti-human interferon-γ (IFN-γ) (B27; BD Pharmingen, San Diego, CA) in endotoxin-free Dulbecco’s-phosphate buffered saline (D-PBS). The plates were then washed three times with D-PBS containing 0.25% Tween-20 (D-PBS/Tween), blocked for 2 hours with RPMI-1640 supplemented with 10% fetal bovine serum, 100U/ml penicillin and 100 μg/ml streptomycin (RPMI-10). Then peptide pool and 2 x 10^5^ PBMCs were added to each well in 100 μl reaction volumes in triplicate. Peptide pool had a total of 236 -15 amino acids overlapping by 11 amino acids peptides that covered the entire sequence of the SIVconsv239 immunogen. Each peptide in a pool was present at a 1 μg/ml concentration. Following an 18-hour incubation at 37°C, the plates were washed nine times with D-PBS/Tween and once with distilled water. The plates were then incubated with 2 μg/ml biotinylated rabbit anti-human IFN-γ (U-Cytech) for 2 hours at room temperature, washed six times with D-PBS/Tween, and incubated for 2.5 hours with a 1:500 dilution of streptavidin-AP (Southern Biotechnology Associates, Birmingham, AL). After five washes with D-PBS/Tween and one with D-PBS, the plates were developed with NBT/BCIP chromogen (Pierce, Rockford, IL), stopped by washing with tap water, air dried, and read with an ELISpot reader (Cellular Technology Ltd., Cleveland, OH). The number of spot forming cells per 10^6^ PBMCs was calculated.

### Activation-induced marker (AIM) and Intracellular cytokine staining (ICS) assay

To perform AIM assay, peripheral blood mononuclear cells (PBMC) were incubated in RPMI-10 at 37°C in a 5% CO_2_ environment for 18 hours in the presence of either DMSO (unstimulated), 1 μg/ml SIVconsv239 peptide pool or 0.4μg/well staphylococcal enterotoxin B (SEB) (Sigma) as positive control. Following stimulation, stimulated cells were then stained with viability dye and a cocktail of antibodies. For ICS assays, peripheral blood mononuclear cells (PBMC) were incubated for 6 hours exactly as described above for AIM assay. All wells contained 2μM of a protein transport inhibitor, monensin (GolgiStop; BD Biosciences), 1μg/ml anti-human CD49d antibody, anti-CD154, anti-CD107a and anti-CD185-biotin. The stimulated cells were then stained with viability dye and the following antibodies to CD4, CD8, CD20, CD45, CD279, CD95, CD28, CD25 and CD14. After fixation and permeabilization with Cytofix/Cytoperm solution (BD Biosciences), the cells were stained with antibodies to CD3, IFNγ, TNFα, IL2, IL- 12, CD69, IL-10, IL-21, IL-17, IL-4, granzyme B, and FoxP3 and fixed with 1% formaldehyde. Antibody details and concentrations are listed in **Table S3**. Samples were collected on a LSR II flow cytometer (BD Biosciences) and analyzed by using FlowJo software (TreeStar). Approximately 250,000 to 900,000 events were collected per sample.

### Binding antibody assay

Enzyme-linked immunosorbent assay (ELISA) were performed to determine the level of binding antibody titers in plasma of immunized monkeys. The assay conducted in 384 well plates (Costar #3700) were coated overnight at 4°C with 2μg/ml of SIVmac251 p55 Gag (BEI Resources, ARP-13384) in 0.1M sodium bicarbonate, then washed (PBS/0.1% Tween-20) and blocked with assay diluent (PBS/4% whey protein/15% Normal Goat Serum/0.5% Tween-20/0.05% sodium azide). Following washing, plasma samples were incubated in a 12 by 3-fold serial dilution starting at 1:30 for 90 minutes. After washing, 10 μl goat anti-monkey IgG-HRP secondary antibody (Jackson ImmunoResearch, #109-035-008)) diluted 1:4,000 in assay diluent without azide was incubated for 1 hour, washed, and followed by detection with 20µl SureBlue Reserve (KPL 53-00-03) for 15 minutes. Reaction was stopped with the addition of 20μl HCL stop solution. Plates were read at 450nm.

### Data analysis

Data were plotted using Prism GraphPad 8.0. All statistical analyses were performed in R version 4.3.1 and Prism 8.0. One-way ANOVA with Bonferroni correction was used to compare multiple independent study groups that with normal distribution. Kruskal-Wallis with or without Dunn’s correction as used when the data had some skewing.

## Supporting information

Supplmentary Materials Santra et al 2026

## Acknowledgements

The authors thank Dr. Nancy Miller of DAIDS/NIAID/NIH; Ying Tam and Christopher Barbosa at Acuitas, George Aoyagi, Ashley Miller, and Jane Fontenot from New Iberia Research Center, University of Louisiana at Lafayette, LA.

## Funding

This study was supported by DAIDS/NIAID SVEU, Duke Consortia for HIV/AIDS Vaccine Development (CHAVD) UM1-AI144371. TH and PB are Jenner Institute Investigators.

## Author Contributions

Conceptualization of the study: BFH, BTK, DW, PB

Study design and making of mRNA and ChAdOx1constructs BTK, TH, DW, EK, GD, BD, JS, M-GA, AP, CH

NHP study and all experiments: LS, WE, RB, BM, GM, HT, ML, SS

NHP immunizations and sample collections SP, FV

Preparation of the manuscript SS, PB, DW, BFH.

## Competing interests

Authors declare no competing interests

## Data, code, and materials availability

All data associated with this study are presented in the paper or the Supplementary materials. The lipid nanoparticle formulations used in this study were developed and provided to Institution by Acuitas Therapeutics Inc. of Vancouver, British Columbia. Acuitas Therapeutics will provide the LNP used in this work under a collaborative research and evaluation agreement to investigators that wish to test them in their studies. The ChAdOx1.SIVconsv239 vaccine is available following a request to TH.

## Notes

### Competing Interest Statement

The authors have declared no competing interest.

## References and Notes

1. P. B. Gilbert et al., Neutralization titer biomarker for antibody-mediated prevention of HIV-1 acquisition. Nat Med 28, 1924–1932 (2022).

2. H. X. Liao et al., Co-evolution of a broadly neutralizing HIV-1 antibody and founder virus. Nature 496, 469–476 (2013).

3. D. Fera et al., Affinity maturation in an HIV broadly neutralizing B-cell lineage through reorientation of variable domains. Proc Natl Acad Sci U S A 111, 10275–10280 (2014).

4. F. Gao et al., Cooperation of B cell lineages in induction of HIV-1-broadly neutralizing antibodies. Cell 158, 481–491 (2014).

5. P. Hraber et al., Prevalence of broadly neutralizing antibody responses during chronic HIV-1 infection. AIDS 28, 163–169 (2014).

6. P. L. Moore, C. Williamson, L. Morris, Virological features associated with the development of broadly neutralizing antibodies to HIV-1. Trends Microbiol 23, 204–211 (2015).

7. R. A. Koup et al., Temporal association of cellular immune responses with the initial control of viremia in primary human immunodeficiency virus type 1 syndrome. J Virol 68, 4650–4655 (1994).

8. J. E. Schmitz et al., Control of viremia in simian immunodeficiency virus infection by CD8+ lymphocytes. Science 283, 857–860 (1999).

9. N. Goonetilleke et al., The first T cell response to transmitted/founder virus contributes to the control of acute viremia in HIV-1 infection. J Exp Med 206, 1253–1272 (2009).

10. P. Borrow, H. Lewicki, B. H. Hahn, G. M. Shaw, M. B. Oldstone, Virus-specific CD8+ cytotoxic T-lymphocyte activity associated with control of viremia in primary human immunodeficiency virus type 1 infection. J Virol 68, 6103–6110 (1994).

11. M. Rolland et al., Broad and Gag-biased HIV-1 epitope repertoires are associated with lower viral loads. PLoS One 3, e1424 (2008).

12. P. Kiepiela et al., CD8+ T-cell responses to different HIV proteins have discordant associations with viral load. Nat Med 13, 46–53 (2007).

13. G. D. Gaiha et al., Structural topology defines protective CD8(+) T cell epitopes in the HIV proteome. Science 364, 480–484 (2019).

14. H. Murakoshi et al., Clinical Control of HIV-1 by Cytotoxic T Cells Specific for Multiple Conserved Epitopes. J Virol 89, 5330–5339 (2015).

15. J. Liu et al., Immune control of an SIV challenge by a T-cell-based vaccine in rhesus monkeys. Nature 457, 87–91 (2009).

16. P. A. Mudd et al., Vaccine-induced CD8+ T cells control AIDS virus replication. Nature 491, 129–133 (2012).

17. J. Bruton, T. Hanke, Exploitation of Unconventional CD8 T-Cell Responses Induced by Engineered Cytomegaloviruses for the Development of an HIV-1 Vaccine. Vaccines (Basel) 13, (2025).

18. S. G. Hansen et al., Profound early control of highly pathogenic SIV by an effector memory T-cell vaccine. Nature 473, 523–527 (2011).

19. S. G. Hansen et al., Immune clearance of highly pathogenic SIV infection. Nature 502, 100–104 (2013).

20. P. S. Arunachalam et al., T cell-inducing vaccine durably prevents mucosal SHIV infection even with lower neutralizing antibody titers. Nat Med 26, 932–940 (2020).

21. R. T. Veenhuis, C. C. Garliss, J. R. Bailey, J. N. Blankson, CD8 Effector T Cells Function Synergistically With Broadly Neutralizing Antibodies to Enhance Suppression of HIV Infection. Front Immunol 12, 708355 (2021).

22. Y. Nishimura et al., Immunotherapy during the acute SHIV infection of macaques confers long-term suppression of viremia. J Exp Med 218, (2021).

23. Y. Nishimura et al., Early antibody therapy can induce long-lasting immunity to SHIV. Nature 543, 559–563 (2017).

24. S. Y. Lim et al., Induction of durable remission by dual immunotherapy in SHIV-infected ART-suppressed macaques. Science 383, 1104–1111 (2024).

25. J. Niessl et al., Combination anti-HIV-1 antibody therapy is associated with increased virus-specific T cell immunity. Nat Med 26, 222–227 (2020).

26. M. Rosas-Umbert et al., Administration of broadly neutralizing anti-HIV-1 antibodies at ART initiation maintains long-term CD8(+) T cell immunity. Nat Commun 13, 6473 (2022).

27. M. J. Peluso et al., Correlates of HIV-1 control after combination immunotherapy. Nature 650, 187–195 (2026).

28. Z. Kiani et al., CD8(+) T cell stemness precedes post-intervention control of HIV viraemia. Nature 650, 196–204 (2026).

29. N. Pardi et al., Nucleoside-modified mRNA vaccines induce potent T follicular helper and germinal center B cell responses. J Exp Med 215, 1571–1588 (2018).

30. C. Li et al., Mechanisms of innate and adaptive immunity to the Pfizer-BioNTech BNT162b2 vaccine. Nat Immunol 23, 543–555 (2022).

31. U. Sahin et al., BNT162b2 vaccine induces neutralizing antibodies and poly-specific T cells in humans. Nature 595, 572–577 (2021).

32. E. Bettini, M. Locci, SARS-CoV-2 mRNA Vaccines: Immunological Mechanism and Beyond. Vaccines (Basel) 9, (2021).

33. E. J. Wherry, D. H. Barouch, T cell immunity to COVID-19 vaccines. Science 377, 821– 822 (2022).

34. C. J. Knudson et al., Lipid-nanoparticle-encapsulated mRNA vaccines induce protective memory CD8 T cells against a lethal viral infection. Mol Ther 29, 2769–2781 (2021).

35. J. Ramos da Silva et al., Single immunizations of self-amplifying or non-replicating mRNA-LNP vaccines control HPV-associated tumors in mice. Sci Transl Med 15, eabn3464 (2023).

36. C. Sittplangkoon et al., mRNA vaccine with unmodified uridine induces robust type I interferon-dependent anti-tumor immunity in a melanoma model. Front Immunol 13, 983000 (2022).

37. H. Danz et al., Synergistic effect of nucleoside modification and ionizable lipid composition on translation and immune responses to mRNA vaccines. NPJ Vaccines 10, 212 (2025).

38. L. A. Rojas et al., Personalized RNA neoantigen vaccines stimulate T cells in pancreatic cancer. Nature 618, 144–150 (2023).

39. B. Ondondo et al., Novel Conserved-region T-cell Mosaic Vaccine With High Global HIV-1 Coverage Is Recognized by Protective Responses in Untreated Infection. Mol Ther 24, 832–842 (2016).

40. M. D. Dicks et al., A novel chimpanzee adenovirus vector with low human seroprevalence: improved systems for vector derivation and comparative immunogenicity. PLoS One 7, e40385 (2012).

41. N. Borthwick et al., Safety and immunogenicity of the ChAdOx1-MVA-vectored conserved mosaic HIVconsvX candidate T-cell vaccines in HIV-CORE 005.2, an open-label, dose-escalation, first-in-human, phase 1 trial in adults living without HIV-1 in the UK. Lancet Microbe 6, 100956 (2025).

42. C. Chanda et al., Safety and broad immunogenicity of HIVconsvX conserved mosaic candidate T-cell vaccines vectored by ChAdOx1 and MVA in HIV-CORE 006: a double-blind, randomised, placebo-controlled phase 1 trial in healthy adults living without HIV-1 in eastern and southern Africa. Lancet Microbe, 101041 (2025).

43. C. L. Gay et al., MVA.HIVconsvX vaccination-evoked T cell expansion inversely associates with age in people with HIV-1 on antiretroviral therapy. J Clin Invest 136, (2026).

44. E. A. Aunins et al., An Il12 mRNA-LNP adjuvant enhances mRNA vaccine-induced CD8 T cell responses. Sci Immunol 10, eads1328 (2025).

45. M. J. Kuroda et al., Analysis of Gag-specific cytotoxic T lymphocytes in simian immunodeficiency virus-infected rhesus monkeys by cell staining with a tetrameric major histocompatibility complex class I-peptide complex. J Exp Med 187, 1373–1381 (1998).

46. B. Brook et al., Adjuvantation of a SARS-CoV-2 mRNA vaccine with controlled tissue-specific expression of an mRNA encoding IL-12p70. Sci Transl Med 16, eadm8451 (2024).

47. N. Agrawal et al., mRNA-based tuberculosis vaccines BNT164a1 and BNT164b1 are immunogenic, well tolerated and efficacious in rodent models. Nat Immunol 27, 1653– 1665 (2026).

48. S. Hervas-Stubbs et al., Direct effects of type I interferons on cells of the immune system. Clin Cancer Res 17, 2619–2627 (2011).

49. K. Kariko, M. Buckstein, H. Ni, D. Weissman, Suppression of RNA recognition by Toll-like receptors: the impact of nucleoside modification and the evolutionary origin of RNA. Immunity 23, 165–175 (2005).

50. R. Verbeke, M. J. Hogan, K. Lore, N. Pardi, Innate immune mechanisms of mRNA vaccines. Immunity 55, 1993–2005 (2022).

51. K. Kariko et al., Incorporation of pseudouridine into mRNA yields superior nonimmunogenic vector with increased translational capacity and biological stability. Mol Ther 16, 1833–1840 (2008).

52. C. Chanda et al., Safety and broad immunogenicity of HIVconsvX conserved mosaic candidate T-cell vaccines vectored by ChAdOx1 and MVA in HIV-CORE 006: a double-blind, randomised, placebo-controlled phase 1 trial in healthy adults living without HIV-1 in eastern and southern Africa. Lancet Microbe 6, 101041 (2025).

53. E. G. Wee et al., Effect of epitope variant co-delivery on the depth of CD8 T cell responses induced by HIV-1 conserved mosaic vaccines. Mol Ther Methods Clin Dev 21, 741–753 (2021).

54. E. G. Wee et al., Parallel Induction of CH505 B Cell Ontogeny-Guided Neutralizing Antibodies and tHIVconsvX Conserved Mosaic-Specific T Cells against HIV-1. Mol Ther Methods Clin Dev 14, 148–160 (2019).

55. B. J. Laidlaw, J. E. Craft, S. M. Kaech, The multifaceted role of CD4(+) T cells in CD8(+) T cell memory. Nat Rev Immunol 16, 102–111 (2016).

56. O. Engstrand et al., Immunogenicity and innate immunity to high-dose and repeated vaccination of modified mRNA versus unmodified mRNA. Mol Ther Nucleic Acids 36, 102588 (2025).

57. M. S. Gebre et al., Optimization of non-coding regions for a non-modified mRNA COVID-19 vaccine. Nature 601, 410–414 (2022).

58. M. Kobayashi et al., Identification and purification of natural killer cell stimulatory factor (NKSF), a cytokine with multiple biologic effects on human lymphocytes. J Exp Med 170, 827–845 (1989).

59. A. S. Stern et al., Purification to homogeneity and partial characterization of cytotoxic lymphocyte maturation factor from human B-lymphoblastoid cells. Proc Natl Acad Sci U S A 87, 6808–6812 (1990).

60. G. Trinchieri, Interleukin-12 and the regulation of innate resistance and adaptive immunity. Nat Rev Immunol 3, 133–146 (2003).

61. C. S. Hsieh et al., Development of TH1 CD4+ T cells through IL-12 produced by Listeria-induced macrophages. Science 260, 547–549 (1993).

62. R. A. Seder, R. Gazzinelli, A. Sher, W. E. Paul, Interleukin 12 acts directly on CD4+ T cells to enhance priming for interferon gamma production and diminishes interleukin 4 inhibition of such priming. Proc Natl Acad Sci U S A 90, 10188–10192 (1993).

63. R. Manetti et al., Natural killer cell stimulatory factor (interleukin 12 [IL-12]) induces T helper type 1 (Th1)-specific immune responses and inhibits the development of IL-4-producing Th cells. J Exp Med 177, 1199–1204 (1993).

64. W. C. Kieper, M. Prlic, C. S. Schmidt, M. F. Mescher, S. C. Jameson, Il-12 enhances CD8 T cell homeostatic expansion. J Immunol 166, 5515–5521 (2001).

65. J. M. Curtsinger, C. M. Johnson, M. F. Mescher, CD8 T cell clonal expansion and development of effector function require prolonged exposure to antigen, costimulation, and signal 3 cytokine. J Immunol 171, 5165–5171 (2003).

66. J. Valenzuela, C. Schmidt, M. Mescher, The roles of IL-12 in providing a third signal for clonal expansion of naive CD8 T cells. J Immunol 169, 6842–6849 (2002).

67. J. Chang et al., IL-12 priming during in vitro antigenic stimulation changes properties of CD8 T cells and increases generation of effector and memory cells. J Immunol 172, 2818–2826 (2004).

68. Z. Xiao, K. A. Casey, S. C. Jameson, J. M. Curtsinger, M. F. Mescher, Programming for CD8 T cell memory development requires IL-12 or type I IFN. J Immunol 182, 2786– 2794 (2009).

69. S. Gurunathan, C. Prussin, D. L. Sacks, R. A. Seder, Vaccine requirements for sustained cellular immunity to an intracellular parasitic infection. Nat Med 4, 1409–1415 (1998).

70. S. A. Calarota, D. B. Weiner, Enhancement of human immunodeficiency virus type 1- DNA vaccine potency through incorporation of T-helper 1 molecular adjuvants. Immunol Rev 199, 84–99 (2004).

71. M. A. Chattergoon et al., Co-immunization with plasmid IL-12 generates a strong T-cell memory response in mice. Vaccine 22, 1744–1750 (2004).

72. P. Wang et al., Re-designing Interleukin-12 to enhance its safety and potential as an anti-tumor immunotherapeutic agent. Nat Commun 8, 1395 (2017).

73. R. A. Elsner, S. Smita, M. J. Shlomchik, IL-12 induces a B cell-intrinsic IL-12/IFNgamma feed-forward loop promoting extrafollicular B cell responses. Nat Immunol 25, 1283–1295 (2024).

74. V. Appay, T. Yamamoto, A. Saez-Cirion, Renaissance of antiviral CD8(+) T cell immunity in vaccination and disease. Nat Rev Immunol 26, 577–590 (2026).

75. L. Stamatatos, The Germline Targeting Vaccine Concept: Overview and Updates from HIV Pre-Clinical and Clinical Trials. Curr HIV Res 23, 380–390 (2026).

76. B. F. Haynes et al., Strategies for HIV-1 vaccines that induce broadly neutralizing antibodies. Nat Rev Immunol 23, 142–158 (2023).

77. D. R. Collins, G. D. Gaiha, B. D. Walker, CD8(+) T cells in HIV control, cure and prevention. Nat Rev Immunol 20, 471–482 (2020).

78. L. A. Jackson et al., An mRNA Vaccine against SARS-CoV-2-Preliminary Report. N Engl J Med 383, 1920–1931 (2020).

79. N. Pardi et al., Zika virus protection by a single low-dose nucleoside-modified mRNA vaccination. Nature 543, 248–251 (2017).

80. N. Pardi et al., Administration of nucleoside-modified mRNA encoding broadly neutralizing antibody protects humanized mice from HIV-1 challenge. Nat Commun 8, 14630 (2017).

81. M. G. Alameh et al., A multivalent mRNA-LNP vaccine protects against Clostridioides difficile infection. Science 386, 69–75 (2024).

82. B. F. Haynes, C. H. Yeh, L. R. Baden, Induction of HIV broadly neutralizing antibodies with vaccination in humans in discovery medicine trials. Curr Opin HIV AIDS, (2026).

83. Z. Mu et al., mRNA-encoded HIV-1 Env trimer ferritin nanoparticles induce monoclonal antibodies that neutralize heterologous HIV-1 isolates in mice. Cell Rep 38, 110514 (2022).

84. E. Melzi et al., Membrane-bound mRNA immunogens lower the threshold to activate HIV Env V2 apex-directed broadly neutralizing B cell precursors in humanized mice. Immunity 55, 2168–2186 e2166 (2022).

85. Z. Mu et al., Comparison of the immunogenicity of mRNA-encoded and protein HIV-1 Env-ferritin nanoparticle designs. J Virol 98, e0013724 (2024).

86. P. Ramezani-Rad et al., Vaccination with an mRNA-encoded membrane-bound HIV envelope trimer induces neutralizing antibodies in animal models. Sci Transl Med 17, eadw0721 (2025).

87. J. R. Willis et al., Vaccination with mRNA-encoded nanoparticles drives early maturation of HIV bnAb precursors in humans. Science 389, eadr8382 (2025).

88. M. G. Alameh et al., Lipid nanoparticles enhance the efficacy of mRNA and protein subunit vaccines by inducing robust T follicular helper cell and humoral responses. Immunity 54, 2877–2892 e2877 (2021).

89. M. Baiersdorfer et al., A Facile Method for the Removal of dsRNA Contaminant from In Vitro-Transcribed mRNA. Mol Ther Nucleic Acids 15, 26–35 (2019).

90. M. A. Maier et al., Biodegradable lipids enabling rapidly eliminated lipid nanoparticles for systemic delivery of RNAi therapeutics. Mol Ther 21, 1570–1578 (2013).

91. M. Jayaraman et al., Maximizing the potency of siRNA lipid nanoparticles for hepatic gene silencing in vivo. Angew Chem Int Ed Engl 51, 8529–8533 (2012).

92. M. D. J. Dicks et al., A novel chimpanzee adenovirus vector with low human seroprevalence: improved systems for vector derivation and comparative immunogenicity. PLoS ONE 7, e40385 (2012).

