## Supplementary material for "Impact of RNA modification and IL-12 boosts on mRNA-LNP vaccine-elicited CD8+ T cell responses": Supplmentary Materials Santra et al 2026

**This file includes:**

Materials and Methods

Figures. S1 to S7

Tables S1 to S3

**Table S1.** Immunization schedule

| Groups (n=5 per group) | Immunizations (IM) |
|---|---|
| 100% 1-methylpseudouridine modified mRNA (50 μg) | Weeks 0, 4, 8 |
| 70% 1-methylpseudouridine with 30% unmodified mRNA (50 μg) | Weeks 0, 4, 8 |
| 30% 1-methylpseudouridine with 70% unmodified mRNA (50 μg) | Weeks 0, 4, 8 |
| 100% unmodified mRNA (50 μg) | Weeks 0, 4, 8 |
| Chimp AdOx1 (5 x$10^{10}$ v.p.) | Weeks 4, 8 |

**Table S2.** Immunization schedule for mRNA-IL-12 boost

| Phase I | Phase II | mRNA/ IL-12 boost (SC) |
|---|---|---|
| 100% modified SIVconsv239-mRNA (50 μg) | same as Phase I + IL-12-mRNA | Weeks 41, 58 |
| 70% modified and 30% unmodified SIVconsv239-mRNA (50 μg) | same as Phase I + IL-12-mRNA | Weeks 41, 58 |
| 30% modified and 70% unmodified SIVconsv239-mRNA (50 μg) | same as Phase I + IL-12-mRNA | Weeks 41, 58 |
| 100% unmodified SIVconsv239-mRNA (50 μg) | same as Phase I + IL-12-mRNA | Weeks 41, 58 |
| ChAdOx1.SIVconsv239 (5×$10^{10}$vp) | 100% modified mRNA (50 μg) + IL-12-mRNA/ | Weeks 41, 58 |

**Figure S1A-C**. **Schema and sequence of mRNA-encoded SIV239 immunogen. (A) SIV gag-pol map depicting viral genome landmarks and polyprotein products**.

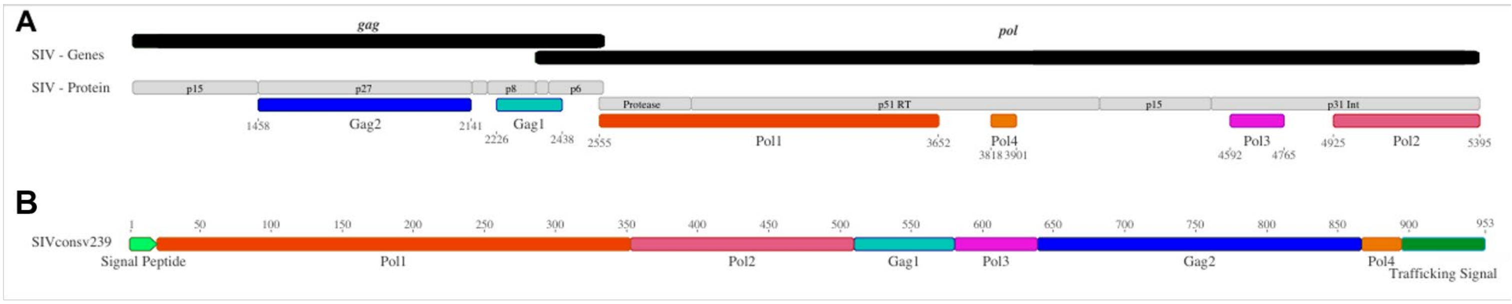


**C**

MRMQLLLLIALSLALVTNSLGPHYTPKIVGGIGGFINTKEYKNVEIEVLGKRIKGTIMTGDTPINIFGRNLLTALGMSLNLPIAKVEPVKVA
LKPGKVGPKLKQWPLSKEKIVALREICEKMEKDGQLEEAPPTNPYNTPTFAIKKKDKNKWRMLIDFRELNRVTQDFTEVQLGIPHPAGL
AKRKRITVLDIGDAYFSIPLDEEFRQYTAFTLPSVNNAEPGKRYIYKVLPQGWKGSPAIFQYTMRHVLEPFRKANPDVTLVQYMDDILIA
SDRTDLEHDRVVLQLKELLNSIGFSTPEEKFQKDPPFQWMGYELWPTKWKLQKIELPQRETWTVNDIQKLVGVLNWAAQIYHTFGVPY
NPQSQGVVEAMNHHLKNQIDRIREQANSVETIVLMAVHCMNFKRRGGIGDMTPAERLINMITTEQEIQFQQSKNSKFKNFRVYYREGR
DQLWKGPGELLWKGEGAVILKVGTDIKVVPRRKAKIIKDYGGGKEVDSSSHMEDTGEAREVAKCWNCGKEGHSAKQCRAPRRQGCW
KCGKMDHVMAKCPDRQAGFLGLGPWGKKPRNFPMAQVHQGLTPTAPPFGLPRLVAKQIVDTCDKCHQKGEAIHGQVNSDLGTWQM
DCTHLEGKIIIVAVHVASGFPVQQIGGNYVHLPLSPRTLNAWVKLIEEKKFGAEVVPGFQALSEGCTPYDINQMLNCVGDHQAAMQIIRD
IINEEAADWDLQHPQPAPQQGQLREPSGSDIAGTTSSVDEQIQWMYRQQNPIPVGNIYRRWIQLGLQKCVRMYNPTNILDVKQGPKEPF
QSYVDRFYKSLRAEQTDAAVKNWMTQTLLIQNANPDCKLVLKGLGVNPTLEEMLTACQGVGGPGQKARLATVIKSQDNQWSYKIHQE
DKILKVGKFASTIPIVGIVAGLAVLAIVVTGAVVAAVMWRRKSSGGKGGSYSQAASSDSAQGSDVSLTA

**Figure S1A-C**. **Schema and sequence of mRNA-encoded SIV239 immunogen. (A) SIV gag-pol map depicting viral genome landmarks and polyprotein products**. Two regions of gag and four regions of pol that correspond to the HIV-1 conserved epitope vaccine HIVconsvX were selected for inclusion in the SIVconsv239 immunogen. Coordinates correspond to nucleotide positions in the SIV reference genome KU955514. **(B) Diagram of SIVconsv239 immunogen coding sequence**. Six regions from A were assembled akin to the HIVconsvX HIV-1 immunogen and flanked by a signal peptide (N-terminus) and an MHC trafficking signal (C-terminus) to yield a 953-aa polyprotein immunogen. **(C)** Amino acid sequence of the SIVconsv239 immunogen, which was expressed from both mRNA and ChAdOx1 vectors.

**Figure S1D**. **Schema of mRNA-encoded IL-12 fusion protein adjuvant.**

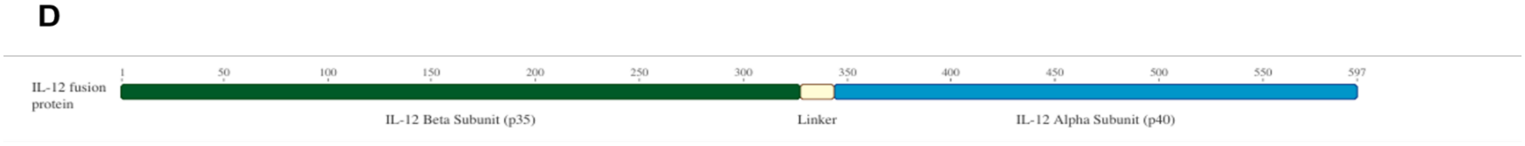


**Figure S1D**. **Schema of mRNA-encoded IL-12 fusion protein adjuvant.** Beta and Alpha subunits were fused with a poly-glycine-serine linker to generate a 597-aa cytokine adjuvant.

**Table S3**

| Reagent | Clone | Amount used/100 µl | Source |
|---|---|---|---|
| Mouse anti-human CD8 FITC | SK1 | 2.5 µl | BD Pharmingen Cat # 347313 |
| Mouse anti-human CD95 CF594 | DX2 | 0.25 µl | BD Pharmingen Cat # 562395 |
| Mouse anti-human CD28 PECy7 | 28.2 | 1.25 µl | LifeTechnologies Cat #25-0289-42 |
| Mouse anti-human CD3 BV421 | SP34.2 | 0.5 µl | BD Pharmingen Cat #562877 |
| Mouse anti-human CD4 BV711 | L200 | 0.75 µl | BD Pharmingen Cat #563913 |
| p11C PE | | 5µl | |
| Mouse anti-human CD196 BB515 | 11A9 | 1.25 µl | BD Pharmingen Cat # 564479 |
| Mouse anti-human CD137 RB613 | 4B4-1 | 0.5 µl | BD Pharmingen Cat #759419 |
| Mouse anti-human CD45RA RB780 | 5H9 | 0.5 µl | BD Pharmingen Cat #755990 |
| Mouse anti-human CD25 RY586 | M-A251 | 4 µl | BD Pharmingen Cat #568459 |
| Mouse anti-human CD69 RY610 | FN50 | 1.25 µl | BD Pharmingen Cat #571167 |
| Hamster anti-human CD278 PE CY7 | C398.4A | 0.25 µl | BioLegend Cat #313520 |
| Mouse anti-human CD194 BV510 | L291H4 | 2.5 µl | BioLegend Cat #359416 |
| Mouse anti-human CD14 BV570 | M5E2 | 1.25 µl | BioLegend Cat #301832 |
| StreptavidinBV650 | | 0.5 µl | BD Pharmingen Cat #563855 |
| Mouse anti-human CD134 BV711 | L106 | 0.5 µl | BD Pharmingen Cat #563855 |
| Mouse anti-human CD4 BV750 | L200 | 0.5 µl | BD Pharmingen Cat #747202 |
| Mouse anti-human CD20 BUV496 | 2H7 | 0.75 µl | BD Pharmingen Cat #569672 |
| Mouse anti-human CD197 BUV563 | 3D12 | 0.8 µl | BD Pharmingen Cat #749679 |
| Mouse anti-NHP CD45 BUV615 | D058-1283 | 0.2 µl | BD Pharmingen Cat #751117 |
| Mouse anti-human CD279 BUV737 | EH12.1 | 2.5 µl | BD Pharmingen Cat #612791 |
| Mouse anti-human CD8 BUV805 | SK1 | 1µl | BD Pharmingen Cat #612889 |
| Mouse anti-human CD3 R718 | SP34.2 | 0.5 µl | BD Pharmingen Cat #566955 |

| | | | |
|---|---|---|---|
| Mouse anti-human CD185 Biotin | MU5UBEE | 2 µl | LifeTechnologies Cat #13-9185-82 |
| Mouse anti-human CD107a BV421 | H4A3 | 2.5 µl | BD Pharmingen Cat #562623 |
| Mouse anti-human CD183 BUV395 | 1C6/CXCR3 | 3µl | BD Pharmingen Cat #565223 |
| LIVE/DEAD™ Fixable Near-IR Dead Cell Stain NIR AMINE | | | Life Technologies Cat # L34976 |
| True-Nuclear™ Transcription Factor Buffer Set | | | BioLegend Cat #424401 |
| Mouse anti-human TNFα FITC | Mab11 | 0.25µl | BD Pharmingen Cat #554512 |
| Mouse anti-human IL12 PE | C8.6 | 0.75µl | LifeTechnologies Cat #12-7129-82 |
| Mouse anti-human CD69 RY610 | FN50 | 1.25µl | BD Pharmingen Cat #571167 |
| Mouse anti-human FoxP3 RY703 | 259D/C7 | 2.5µl | BD Pharmingen Cat #571433 |
| Rat anti-human IL-10 BV421 | JES3-9D7 | 2.5µl | BioLegend Cat #501422 |
| Mouse anti-human Granzyme B BV510 | GB11 | 1.0µl | BD Pharmingen Cat #563388 |
| Mouse anti-human IL-17 BV605 | BL168 | 3.0µl | BioLegend Cat #512326 |
| Rat anti-human IL-4 BV711 | MP4-25D2 | 2.5µl | BD Pharmingen Cat #564112 |
| Mouse anti-human IFN-γ BUV395 | B27 | 2.5µl | BD Pharmingen Cat #563563 |
| Rat anti-human IL-2 BUV737 | MQ1-17H12 | 2.5µl | BD Pharmingen Cat #612836 |
| Mouse anti-human IL-21 A647 | 3A3-N2 | 2.5µl | BioLegend Cat #513006 |
| Mouse anti-human CD154 BV785 | 24-31 | 2.5µl | BioLegend Cat #310842 |
| Mouse anti-human CD107a PE CY5 | H4A3 | 8.5µl | BD Pharmingen Cat #555802 |
| Mouse anti-human CD279 RB705 | EH12.1 | 2.0µl | BD Pharmingen Cat #612791 |
| Mouse anti-human CD95 RB780 | DX2 | 2.5µl | BD Pharmingen Cat #569126 |
| Mouse anti-human CD28 BUV563 | 28.2 | 0.8µl | BD Pharmingen Cat #741392 |

**Figure S2 Gating strategy for tetramer-positive CD8$^+$ T cells in whole blood.**

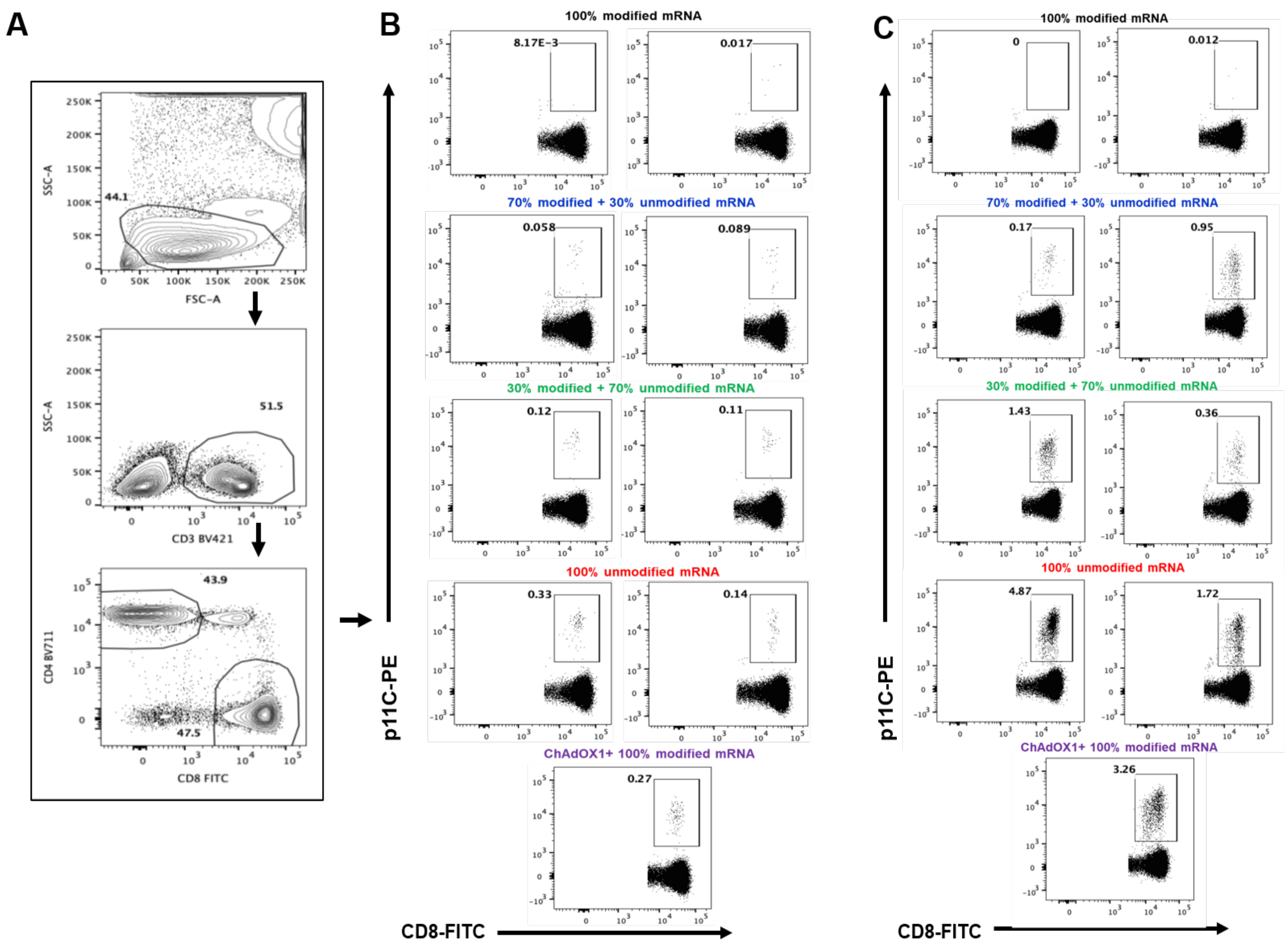


**Figure S2 Gating strategy for tetramer-positive CD8$^+$ T cells in whole blood.** Anti-coagulated whole blood was stained with p11c tetramer and anti-CD3, anti-CD8 antibodies. Gating strategy to determine CD8+p11C+ cells is shown. **A**. Serial gating to define lymphocytes, CD3$^+$ T cells and CD4$^+$ or CD8$^+$ T cells. **B and C**. Representative examples of p11C$^+$ CD8$^+$ T cells in individual *Mamu-A*01*$^+$ monkeys at Study week 12 **(B)** and 59 **(C)**

**Figure S3A. Representative flow cytometry plots illustrating gating strategy for cytokines and degranulation molecules in PBMC.**

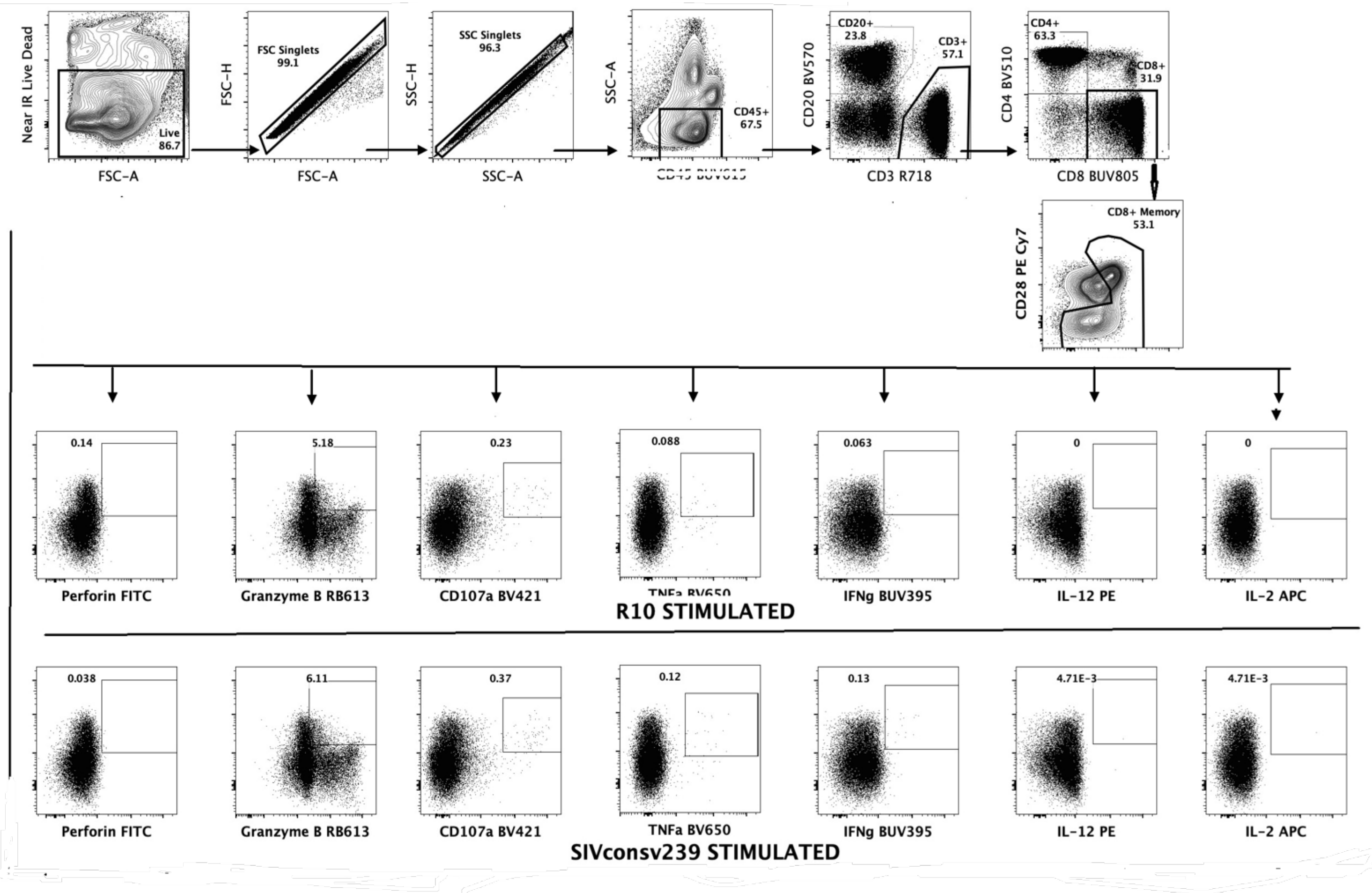


**Figure S3A. Representative flow cytometry plots illustrating gating strategy for cytokines and degranulation molecules in PBMC.** Density plots showing gating of different viable cell populations in black boxes (top) and gating for lytic proteins, the degranulation marker CD107a, and cytokines (rectangles) in both unstimulated (R10) and SIVconsv239 peptide stimulated PBMC. Representative of ten experiments.

**Figure S3B. Representative flow cytometry plots illustrating gating strategy for Activation-Induced Marker (AIM) assay.**

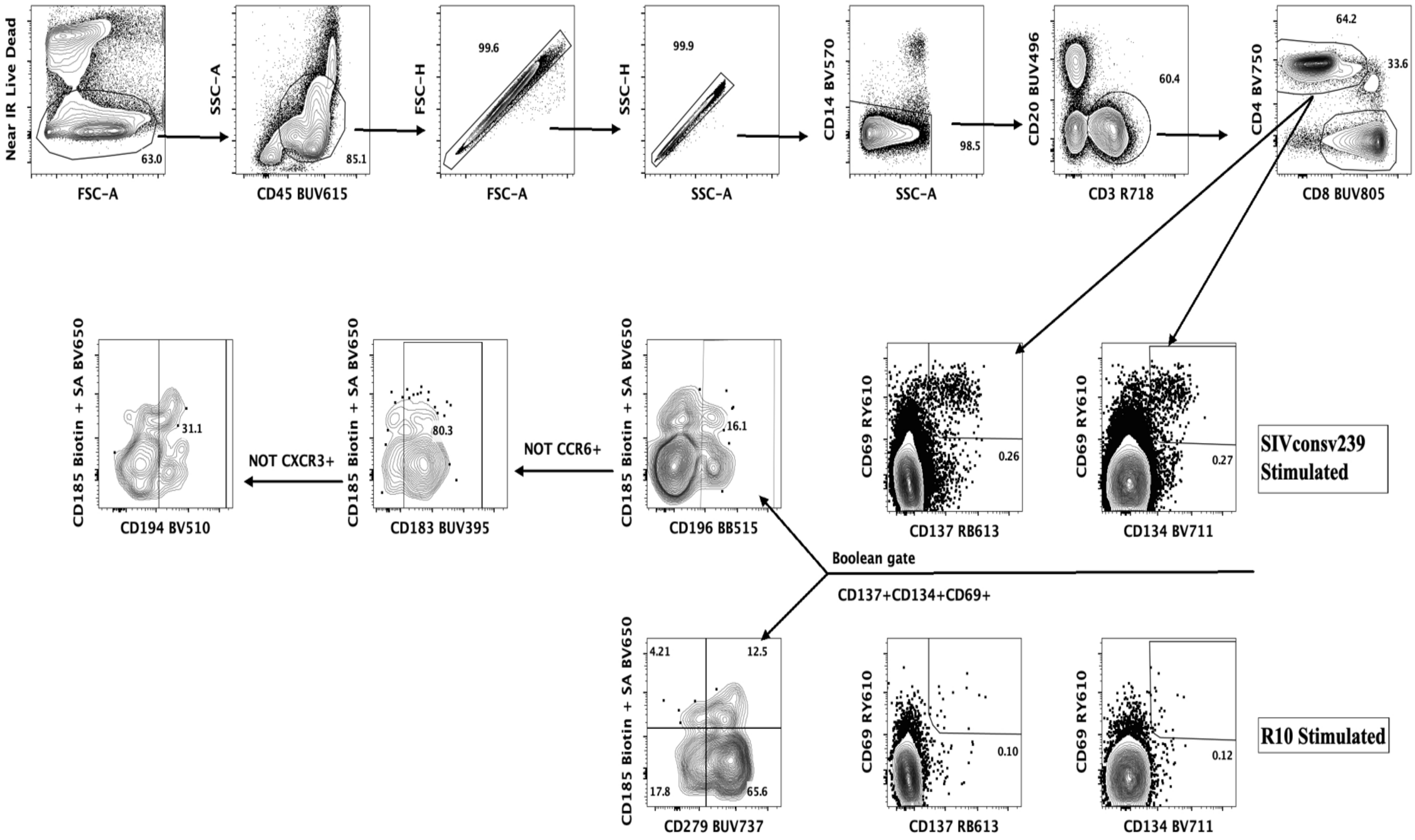


**Figure S3B. Representative flow cytometry plots illustrating gating strategy for AIM assay.** Density plots showing gating of different viable cell populations in black boxes, circles and rectangles in both unstimulated (R10) and SIVconsv239 peptide stimulated PBMC. Representative of four experiments.

**Figure S4. $T_{H2}$-type cytokine secretion profiles of CD4$^+$ (A) and CD8$^+$ (B) T lymphocytes of the mRNA and ChAdOX1-vaccinated monkeys.**

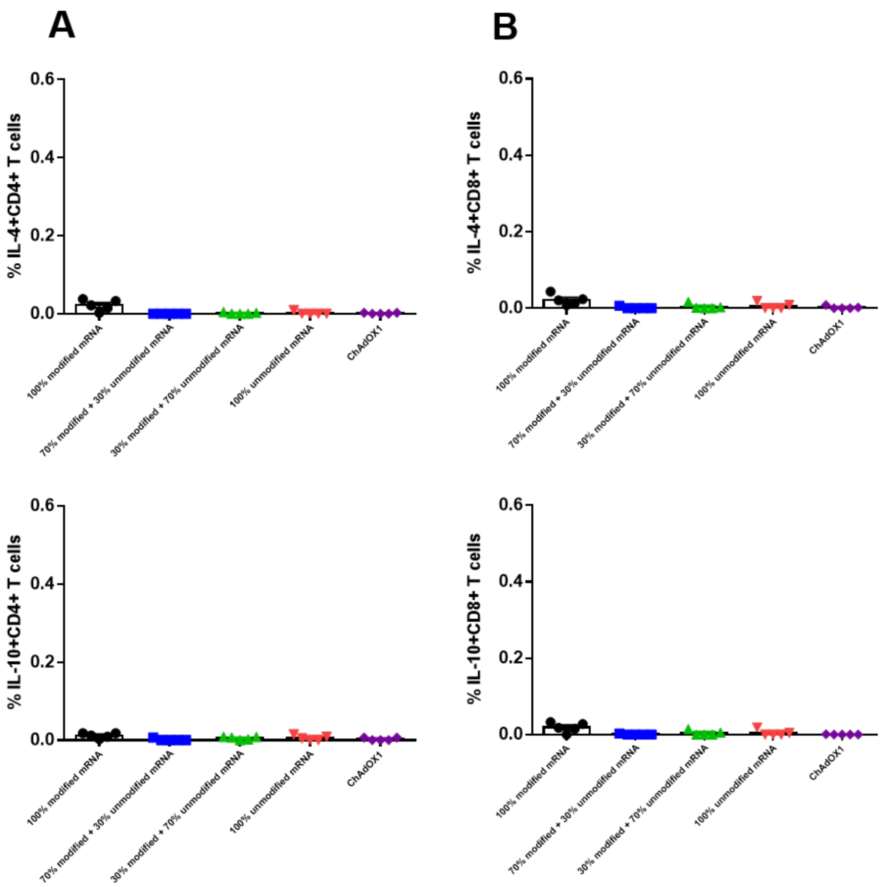


**Figure S4. $T_{H2}$-type cytokine secretion profiles of CD4$^+$ (A) and CD8$^+$ (B) T lymphocytes of the mRNA and ChAdOX1-vaccinated monkeys.** Percent IL-4 (top panels) and IL10 (bottom panels) positive CD4$^+$ and CD8$^+$ T lymphocytes from individual monkeys in each group following stimulation with SIVconsv239 peptide pool are shown on Study week 12- 8 weeks post ChAdOX1 and 4 weeks post final mRNA immunizations following stimulation with SIVconsv239 peptide pool are shown. Mean $\pm$ SEM per group are shown.

**Figure S5. Fold increase in magnitude of ELISpot responses in groups of monkeys post boost with IL-12 encoding mRNA**.

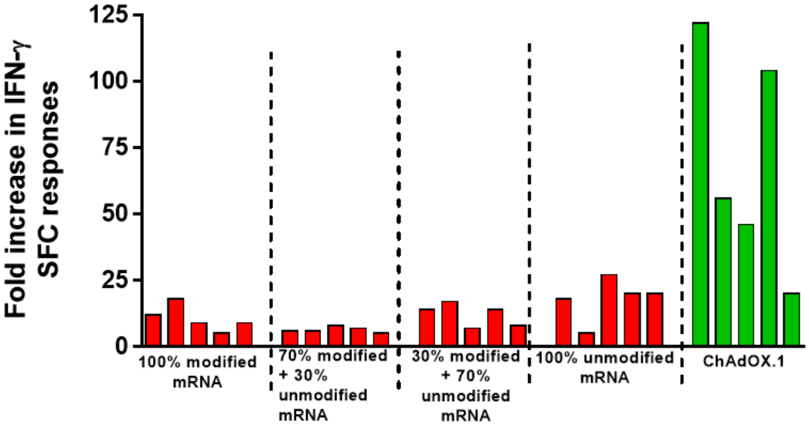


**Figure S5. Fold increase in magnitude of ELISpot responses in groups of monkeys post boost with IL-12 encoding mRNA**. Fold increases in IFN-γ ELISpot response from study week 34 (pre-IL-12 mRNA) to study week 60 (post-IL-12 mRNA) per monkey are shown**.**

**Figure S6. T lymphocytes from the mRNA and ChAdOX1-vaccinated monkeys show cytotoxic potential**.

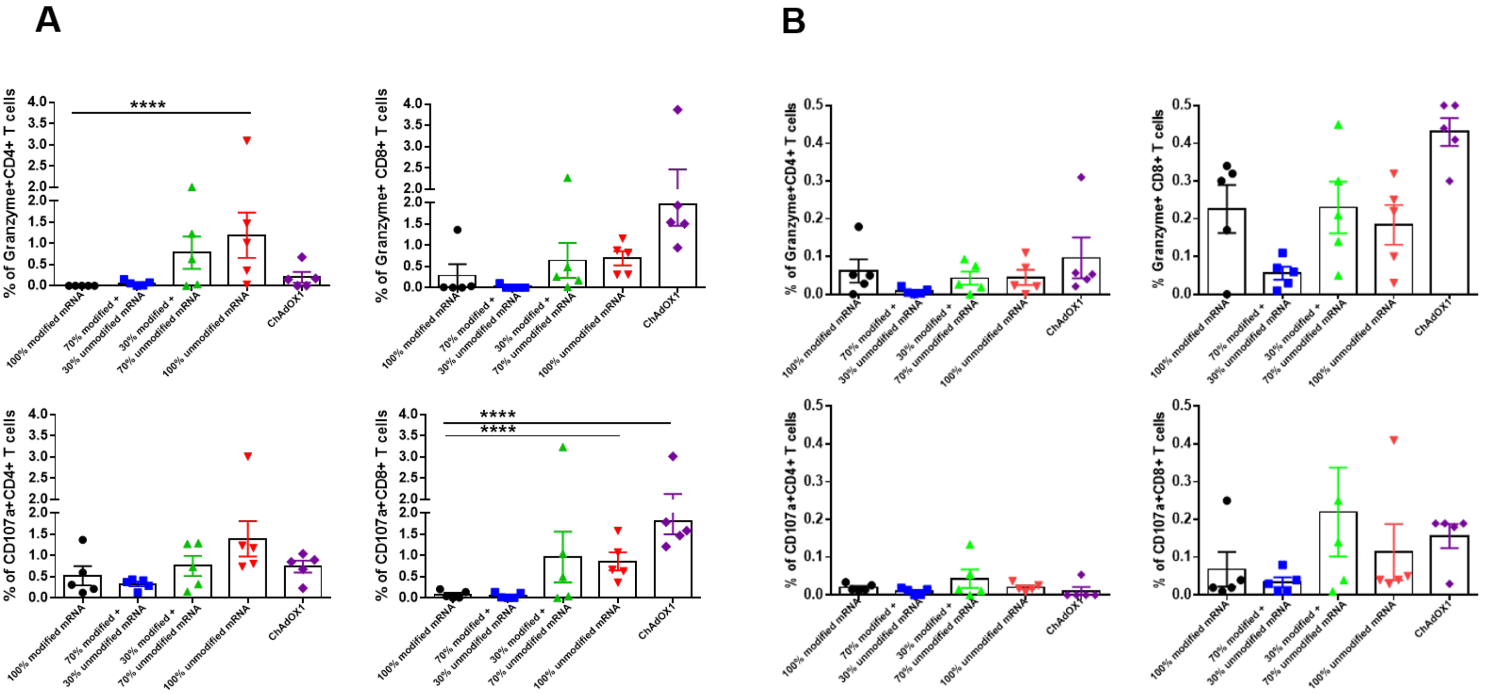


**Figure S6. T lymphocytes from the mRNA and ChAdOX1-vaccinated monkeys show cytotoxic potential**. Percent granzyme B (top panels), and CD107a (bottom panels) positive $CD4^+$ and $CD8^+$ T lymphocytes from individual monkeys in each group following stimulation with SIVconsv239 peptide pool on weeks 59 **(A)** and 78 (**B**) of the study are shown. Mean $\pm$ SEM per group are shown (**** p<0.0001 one-way ANOVA with Bonferroni correction).

**Figure S7. Cytokine expression profile of lymph node CD4$^+$ (top panels) and CD8$^+$ T cells.**

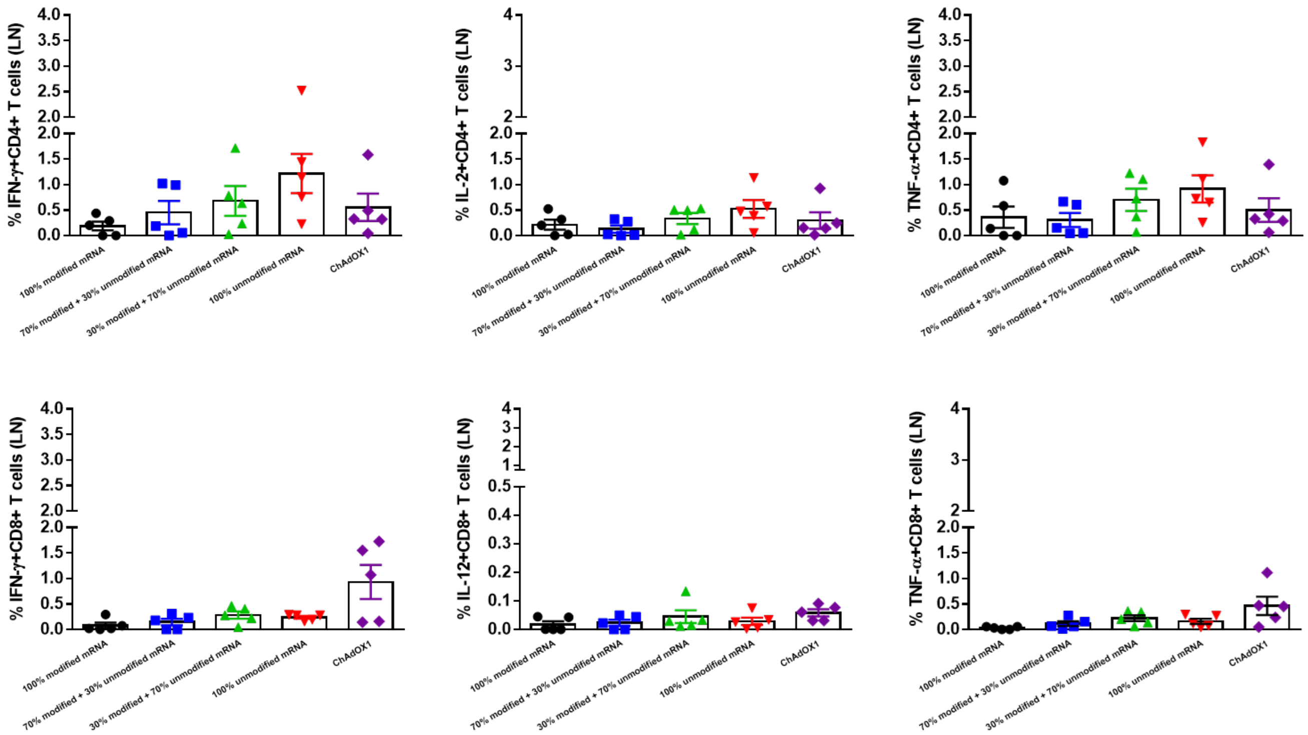


**Figure S7. Cytokine expression profile of lymph node CD4$^+$ (top panels) and CD8$^+$ T cells.** Cytokine expression profile of lymph node CD4$^+$ (top panels) and CD8$^+$ T cells. (bottom panels) one week post final mRNA-IL-12 immunization at week 59 from individual monkeys in each group following stimulation with SIVconsv239 peptide pool. Mean $\pm$ SEM per group are shown.
